# Injector-based pump–probe serial synchrotron crystallography of CrPhotLOV1 at the MicroMAX beamline

**DOI:** 10.64898/2026.10.26.762041

**Authors:** Bence Olasz, Magdalena Wisniewska, Anna Koczurowska, Cecilia Maria Casadei, Aleksandra Solecka, Sofia M. Kapetanaki, Magdalena Pachota, Larissa Lopes Cavalcante, Ayesha Asghar, Dean Lang, Mirko Milas, Thomas Ursby, Madan Kumar Shankar, Manoop Chenchiliyan, Jie Nan, Przemyslaw Nogly

**Author notes:** These authors contributed equally to this work (Bence Olasz, Magdalena Wisniewska).

## Abstract

Time-resolved serial synchrotron crystallography (TR-SSX) has become a powerful approach for investigating structural dynamics in proteins under near-physiological conditions. Its successful application, however, depends on robust sample delivery, reliable pump–probe synchronization, and reproducible data-processing strategies. Here, an injector-based pump–probe TR-SSX workflow at the MicroMAX beamline is applied to investigate the structural dynamics of the LOV1 domain of the blue-light photoreceptor phototropin from *Chlamydomonas reinhardtii* (CrPhotLOV1).

Room-temperature serial datasets were collected for an independently measured laser-off dark state together with four light-activated pump–probe time windows (0–350, 500–850, 1000–1350 and 1500–1850 µs after a microsecond laser pulse). All four time windows and an interleaved pump–probe dark reference were recorded in a single run with the JUNGFRAU detector at 2 kHz. Because thioether adduct formation in CrPhotLOV1 is complete within about 20 µs and the adduct persists for hundreds of seconds, the four delays probe the same photoadduct state and serve as independent replicates. The workflow was evaluated in terms of data quality, light contamination in the pump–probe dark reference, difference Fourier analysis, extrapolated electron-density maps and refinement of light-state models. Consistent activation levels were obtained across all time points, enabling reliable structural interpretation of the light-activated state.

Difference Fourier maps and refined extrapolated models reproducibly revealed formation of the characteristic FMN–Cys57 thioether adduct together with localized rearrangements within the chromophore-binding pocket. The refined light-state structures agree closely with previously reported CrPhotLOV1 models.

These results establish a robust and reproducible workflow for injector-based pump–probe TR-SSX at the MicroMAX beamline, suitable for routine time-resolved studies of light-responsive proteins.

**Synopsis:** An injector-based pump–probe time-resolved serial synchrotron crystallography workflow at the MicroMAX beamline of MAX IV is applied to investigate the structural dynamics of the LOV1 domain of *Chlamydomonas reinhardtii* phototropin: consistent photoadduct structures in four time windows spanning 0–1850 µs, including a 0–350 µs window, with dark references showing no detectable light contamination.

## 1. Introduction

Serial crystallography (SX) merges diffraction data from many microcrystals, so that the X-ray dose is spread over the sample and data can be collected at room temperature (Chapman *et al*., 2014; Chapman, 2017; Mehrabi *et al*., 2021), although radiation damage can still occur, even with femtosecond X-ray pulses (Lomb *et al*., 2011; Nass *et al*., 2015). Because reactions can be initiated rapidly and uniformly in small crystals, for example by light activation of photosensitive proteins, SX is also well suited to time-resolved crystallography (TRX), which resolves transient structural states and thereby the dynamics of biological processes (Moffat, 1989, 2001; Schmidt, 2008, 2013, 2020, 2023; Pearson & Mehrabi, 2020).

Time resolved macromolecular crystallography (TRX) was pioneered at synchrotrons using Laue diffraction from single crystals (Moffat, 1989; Genick *et al*., 1997; Schotte *et al*., 2012). The serial approach, in which each crystal is measured only once, was developed at X-ray free electron laser (XFEL) facilities and subsequently transferred to synchrotron beamlines as serial synchrotron crystallography (SSX), which in combination with reaction triggering, such as by laser, enables time resolved SSX (TR-SSX). TRX has been successfully applied to a broad spectrum of light sensitive proteins including light driven proton pump bacteriorhodopsin, sensory rhodopsin II, as well as photoactive yellow protein and LOV domain containing proteins, demonstrating its capability to resolve light induced conformational changes at physiologically relevant temperatures (Weinert *et al*., 2019; Bosman *et al*., 2025; Genick *et al*., 1997; Ren *et al*., 2001; Ihee *et al*., 2005; Schotte *et al*., 2012; Jung *et al*., 2013; Gotthard, Mous *et al*., 2024; Aumonier *et al*., 2020; Malla *et al*., 2025). Notably, a systematic head-to-head comparison of crystallographic diffraction data collected using XFEL based serial femtosecond crystallography (SFX) and SSX, performed on identical crystals under matched experimental conditions, demonstrated that both approaches yield diffraction data of comparable quality (Mehrabi *et al*., 2021). This study demonstrated that data quality is governed primarily by crystal properties rather than the radiation source, underscoring that SSX is capable of supporting time resolved studies using a radiation source selected to match the required time resolution.

Time-resolved serial measurements at synchrotrons became practical with the higher flux density of third-and, more recently, fourth-generation sources and with beamlines that combine microfocus X-ray beams with fast, low-noise detectors (Pearson & Mehrabi, 2020; Martin-Garcia, 2021; Nam, 2024), now operating at kilohertz frame rates at fourth-generation sources (Leonarski *et al*., 2023), and with X-ray choppers that shorten exposures to the microsecond range (Gonzalez *et al*., 2025). As a result, SSX is now available at microfocus beamlines at many high brilliance synchrotron facilities, where beamtime is considerably more accessible than XFEL beamtime (Nam, 2024). Substantial effort has also been devoted to sample delivery, which plays a critical role in SSX experiments. Injector-based SSX with viscous jets was established early (Nogly *et al*., 2015; Botha *et al*., 2015) and subsequently extended to time resolved studies with 5 ms frames (Weinert *et al*., 2019) and, at the P14 beamline of PETRA III, to a single sub-millisecond snapshot of the xenorhodopsin BcXeR, in which a 500 µs X-ray exposure was delayed by 250 µs relative to a 500 µs pump pulse (Kovalev *et al*., 2023), while fixed-target approaches such as the hit-and-return (HARE) strategy reduce sample consumption, provide high hit rates and support time resolved photoactivation and mixing experiments (Schulz *et al*., 2018; Mehrabi *et al*., 2019, 2020). Collectively, these developments have established SSX as an approach for room temperature structure studies and time-resolved investigations of protein dynamics while limiting radiation damage.

MicroMAX is a fourth-generation macromolecular crystallography beamline with dedicated capabilities for (time-resolved) serial crystallography, bridging synchrotron and XFEL capabilities, combining high X-ray flux, tunable beam sizes, a fast integrating detector and advanced time resolved instrumentation (Gonzalez *et al*., 2025, 2026). A multilayer monochromator provides a high-flux X-ray beam that can be focused to 10 (H) × 5 (V) µm², with approximately 10¹⁴ photons s⁻¹ at the sample position, while an X-ray chopper enables control of the X-ray pulse duration down to 10 µs. Fast diffraction data acquisition is achieved by a JUNGFRAU 9M integrating detector (Mozzanica *et al*., 2018; Leonarski *et al*., 2023). A tunable nanosecond laser system and several laser diodes operating at different wavelengths are available for triggering light-induced reactions, depending on the proteins. In combination with the high viscosity extruder (HVE) sample delivery, these components can be orchestrated and synchronized through a PandABox timing and control system (Zhang *et al*., 2018; Gonzalez *et al*., 2025), allowing the experimental scheme to be readily adapted to the requirements of individual experiments. Combined with automated data analysis pipelines, this flexible instrumentation makes MicroMAX well suited for time resolved crystallography on protein microcrystals.

A central challenge in TRX experiments is the precise and synchronous initiation of biochemical reactions within crystalline samples. In this context, blue-light photoreceptors such as photoactive yellow protein (PYP) and LOV domains have become widely used systems (Genick *et al*., 1997; Ren *et al*., 2001; Ihee *et al*., 2005; Schotte *et al*., 2012; Jung *et al*., 2013; Gotthard, Mous *et al*., 2024; Aumonier *et al*., 2020; Malla *et al*., 2025). Their intrinsic light sensitivity enables rapid, homogeneous activation via laser excitation, eliminating the need for diffusive mixing and allowing well-defined pump–probe experiments. These properties make photoactive proteins particularly valuable for methodological development and benchmarking, and for investigating structural mechanisms underlying signal transduction. Importantly, such systems also provide a critical link between synchrotron and XFEL-based TRX, enabling the integration of structural snapshots from femtosecond to millisecond timescales into coherent reaction pathways. As a result, photoactive proteins continue to play a central role in advancing both the methodological capabilities and the biological impact of time-resolved crystallography.

Light-Oxygen-Voltage (LOV) domains are small flavin-binding sensory modules that convert blue light into intracellular signals (Salomon *et al*., 2000; Crosson & Moffat, 2001). Upon blue-light absorption, the flavin mononucleotide (FMN) chromophore is excited to a triplet state, which decays into a covalent thioether adduct between the FMN C4a atom and a conserved reactive cysteine (Swartz *et al*., 2001; Kottke *et al*., 2003). Adduct formation triggers intramolecular signaling that ultimately modulates the activity of downstream effector domains (Harper *et al*., 2003). In phototropin Phot-1 from *Chlamydomonas reinhardtii*, two LOV domains, LOV1 and LOV2, are followed by a Ser/Thr kinase effector domain, and the conformation of each linker that couples a LOV domain to the kinase changes with the signaling state of that domain (Huang *et al*., 2002; Okajima *et al*., 2014; Henry *et al*., 2020). In the isolated LOV1 domain (CrPhotLOV1), the triplet state decays with time constants of 0.8 and 4 µs into the adduct state, which reverts to the dark state within hundreds of seconds (Kottke *et al*., 2003). Adduct formation is therefore complete on the microsecond timescale, and any delay between a few microseconds and several seconds probes the same long-lived photoadduct state. CrPhotLOV1 has been characterized structurally by cryo-crystallography (Fedorov *et al*., 2003), by TR-SSX at millisecond delays (Gotthard, Mous *et al*., 2024) and as a test system for fixed-target pump–probe SFX at the SwissFEL Cristallina end station (Gotthard, Flores-Ibarra *et al*., 2024), and most recently by TR-SFX at picosecond-to-microsecond delays, which captured the formation of the covalent FMN–Cys57 bond (Olasz *et al*., 2026). Because they act as natural light-controlled switches, LOV domains are also widely used as building blocks of optogenetic tools (Losi *et al*., 2018).

In this report, we extend our previous TR-SSX work on CrPhotLOV1 at millisecond delays (Gotthard, Mous *et al*., 2024) to the microsecond regime at the MicroMAX beamline of the fourth-generation MAX IV synchrotron, using the same microcrystal preparation and injector-based delivery, and we describe in detail the pump–probe data collection and processing. Using this approach, we acquired datasets in four pump–probe time windows (0–350, 500–850, 1000–1350 and 1500–1850 µs), enabling reliable characterization of light induced changes. The 0–350 µs window is shorter and ends earlier than the 250–750 µs snapshot obtained for BcXeR with an injector at PETRA III (Kovalev *et al*., 2023). Given the photocycle kinetics of CrPhotLOV1, these delays all probe the fully formed photoadduct state, so the four datasets serve as independent replicates that allow the reproducibility of the workflow, including excitation efficiency and light-contamination control, to be assessed. We demonstrate the absence of light contamination in the pump–probe reference dataset, reproducible formation of the characteristic cysteinyl–FMN photoadduct, and reliable refinement of light-state structural models. The resulting dark-and light-state structures are consistent with previously reported CrPhotLOV1 models (Fedorov *et al*., 2003; Gotthard, Mous *et al*., 2024). Together, these results demonstrate the suitability of MicroMAX and the presented workflow for routine injector-based TR-SSX experiments on the hundreds-of-microseconds timescale.

## 2. Materials and methods

### 2.1. Expression and purification

The genetic sequence encoding amino acids 16–133 of the LOV1 domain of the *Chlamydomonas reinhardtii* phot1 protein was cloned into the pET16b expression vector between the NdeI and XhoI restriction sites, enabling expression of the protein with an N-terminal His-tag, as described previously (Gotthard, Mous *et al*., 2024). Protein expression was carried out in *Escherichia coli* BL21 (DE3) cells grown in ZYP-5052 auto-inducing medium (Studier, 2005). Starter cultures were prepared one day prior to expression by inoculating LB medium supplemented with 100 µg/ml ampicillin either directly with material from a glycerol stock or by picking a single colony from a freshly streaked plate. These cultures were grown overnight at 37 °C with shaking at 160–180 rpm. The following day, production cultures were inoculated with the overnight starter culture (1:125) into LB medium containing 100 µg/ml ampicillin and incubated at 37 °C with shaking until an OD₆ of approximately 1.0 was reached. Protein expression was induced by reducing the temperature to 17 °C and continuing incubation overnight. Cells were harvested by centrifugation (20 min, 4,500 × g, 4 °C), and cell pellets were stored at –80 °C until further processing. Frozen pellets were thawed and resuspended in lysis buffer consisting of 50 mM Tris-HCl pH 8.0, 300 mM NaCl, 20 mM imidazole, and 1 mM MgCl₂, supplemented with DNase I (5 µl per 100 ml), lysozyme (0.5 mg/ml), and one protease inhibitor tablet per 100 ml. Cells were disrupted by a single passage through a continuous flow cell disruptor (Constant Systems, model BT40/TS2/AA) at 24 kPSI. The lysate was clarified by centrifugation (45 min, 110,000 × g, 4 °C) and subsequently incubated with 1 mM FMN for 1 h at 4 °C. The clarified lysate was applied to a pre-equilibrated Ni-NTA affinity column, and bound protein was eluted over a linear imidazole gradient ranging from 20 to 500 mM in 50 mM Tris-HCl pH 8.0, 300 mM NaCl, and 1 mM MgCl₂. Absorbance at 280, 450, and 260 nm was measured, and the 280/450 nm ratio was used to identify protein containing peak fractions. These fractions were then pooled and concentrated to approximately 30 ml by centrifugal ultrafiltration. The concentrated sample was then subjected to desalting on a pre-equilibrated column using the standard protocol, and fractions containing the protein of interest were combined based on their absorbance profiles. The final pooled material was quantified spectrophotometrically, aliquoted, flash frozen in liquid nitrogen, and stored at –80 °C until use.

### 2.2. Crystallization

Limited proteolysis with trypsin removed the purification tag from the purified protein (adding 1:10 of 0.25 mg ml^−1^ trypsin solution; trypsin:protein). Trypsin was added to the protein (at 8.0 mg ml^-1^) prior to setting up the crystallization experiment. Proteolysis was performed in the crystallization setup at 20 °C (proteolysis was unquenched throughout crystallization). The condition producing a high density of microcrystals suitable for serial crystallography has been identified before (Gotthard, Mous *et al*., 2024) and consists of 100 mM sodium cacodylate at pH 6.5 and 1.0 M sodium citrate dibasic trihydrate. Crystallization and improving crystal size homogeneity were achieved in the batch crystallization method with seeding. Crystals obtained during the first round of crystallization were used to prepare a seeding stock by crushing them with seeding beads (Hampton Research) and serially diluted to acquire seeding stocks used during crystallization. The seeds were mixed with protein/trypsin mixture (at 1:10 seed:protein ratio). Finally, the mix was added dropwise in 15 ml Falcon tubes in the above-mentioned crystallization condition in a 2:1 ratio at 20 °C (protein:precipitant, v/v). Crystals with a size of 10–20 µm (Fig. S1) appeared the next day and slowly sedimented at the bottom of the Falcon tube.

### 2.3. Sample preparation for serial synchrotron crystallography

A jetting solution of hydroxyethyl cellulose [23%(w/v)] (Sugahara *et al*., 2017), as used previously for CrPhotLOV1 (Gotthard, Mous *et al*., 2024), was prepared by dissolving dried cellulose in a solution containing the protein purification buffer and the crystallization condition in a 1:2 ratio (purification buffer: crystallization condition). The cellulose mix was left to hydrate at RT until the medium became clear. Crystals were sedimented by centrifugation (800 x g for 1 min) and resuspended in the mother liquor for stabilization at the desired concentration. Resuspended crystals were inserted from the back of a Hamilton syringe and mixed in a 1:1 ratio with the hydrated viscous matrix using a three-way syringe coupler (James *et al*., 2019).

### 2.4. TR-SSX data collection at MicroMAX

A serial time-resolved pump–probe crystallography experiment was conducted at the MicroMAX end station (Fig. 1). The experiment was performed at an X-ray energy of 13.7 keV using the multilayer monochromator (MLM) with a relative energy bandwidth of 1%. The X-ray beam was focused to 20 (H) × 5 (V) µm² FWHM at the sample position using compound refractive lenses (CRLs), with a photon flux of 8.6 × 10¹s photons s⁻¹. An X-ray chopper (Celeroton Ltd.) was operated at 2 kHz to produce 350 µs X-ray pulses. For both dark and time-resolved data collection, diffraction images were recorded with a JUNGFRAU 9M detector (Mozzanica *et al*., 2018), also operated at 2 kHz (500 µs frame time), with a 350 µs integration time synchronized to the X-ray pulses. The dose per X-ray exposure was estimated using RADDOSE-3D (Zeldin *et al*., 2013) to be 25 kGy (average diffraction-weighted dose).

**Figure 1.**
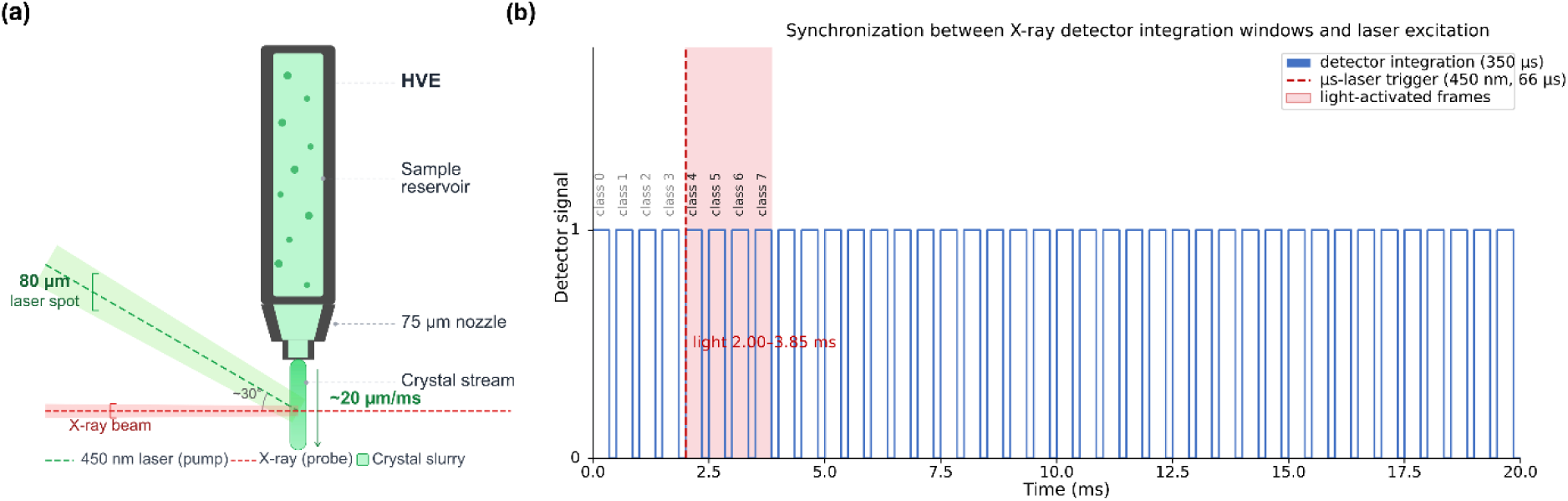
Pump–probe TR-SSX setup at the MicroMAX end station. (*a*) Sample–beam interaction region. The high-viscosity extruder (HVE; 75 µm nozzle) delivers the crystal stream (pale green, ∼20 µm ms^−1^) into the 80 µm spot of the 450 nm pump laser (bright green), which is centered on the X-ray beam (red) and incident at about 30° to it. ( *b*) Timing scheme. Detector frames are recorded at 2 kHz (500 µs period) with 350 µs integration (blue bars). The 66 µs laser pulse is triggered at 2.00 ms (dashed red line) at 50 Hz, so the pattern repeats every 20 ms. Frames of classes 0–3 precede the pulse and form the dark reference; frames of classes 4–7 cover the time windows 0–350, 500–850, 1000–1350 and 1500–1850 µs after excitation (shaded region, 2.00–3.85 ms). Later frames in each cycle were not used.

The sample was delivered using an HVE (Weierstall *et al*., 2014; Botha *et al*., 2015; Shilova *et al*., 2020) equipped with a 75 µm nozzle. A flow rate of 5.32 µL/min produced a stable continuous stream with an estimated linear velocity of approximately 20 µm/ms.

For the pump–probe experiment, photoexcitation was provided by a 450 nm CW laser diode (L450G2, Thorlabs) operated in analog modulation mode. The divergent diode output was collimated with an aspheric lens (C280TMD-A, Thorlabs) and focused with a 100 mm focal-length achromatic doublet (AC254-100-A, Thorlabs) to 86 (H) × 80 (V) µm² (1/e²) at the sample, centered on the X-ray beam and incident at an angle of approximately 30° relative to the X-ray beam axis. Excitation pulses of 66 µs were delivered at 50 Hz, with a pulse energy of 3.3 µJ, corresponding to a mean fluence of 61 mJ cm⁻² averaged over the 1/e² beam area.

Laser excitation and X-ray acquisition were synchronized using a PandABox timing system (Zhang *et al*., 2018). Four light-activated frames were collected over the time windows 0–350, 500–850, 1000–1350 and 1500–1850 µs following excitation, corresponding to midpoints 175, 675, 1175 and 1675 µs (Fig. 1*b*). The PandABox also generated the programmable signal that modulated the laser-diode output, and each laser pulse was synchronized with the start of a detector frame. Throughout the text, datasets are referred to by their respective time windows.

At a flow speed of 20 µm/ms, the sample traveled 37 µm by the end of the last time window (1850 µs), remaining within the laser illuminated region. Between laser pulses at 50 Hz, the stream advances by approximately 400 µm, ensuring each laser pulse excited fresh sample. Frames recorded after the fourth light class, during which the illuminated volume may still pass the X-ray interaction point, were excluded from the analysis.

### 2.5. Data processing

Data were processed using the MicroMAX in-house developed, CrystFEL-based (White *et al*., 2012, 2016) data reduction workflow, which includes automated beam position and peakfinder8 parameter optimisation as well as time-delay classification of frames. CrystFEL version 0.11.1 was used, and images corresponding to each time delay in the activation sequence (class0–class7) were binned as described previously (Gotthard, Mous *et al*., 2024). Indexing and integration were performed with indexamajig using the XGANDALF (Gevorkov *et al*., 2019) algorithm, searching for peaks with peakfinder8 (minimum signal-to-noise of 4.0, threshold 10.0, minimum pixel count of 2.0, local background radius of 3 and minimum number of peaks of 10.0) and using the unit-cell parameters a = b = 122.6 Å, c = 46.6 Å, α = β = 90°, γ = 120°. Peak intensities were integrated using the rings method with integration radii of 2, 5 and 7 pixels. Data were merged and scaled with partialator using the unity partiality model. The resulting hkl files were converted into mtz format with f2mtz from the CCP4 suite (Winn *et al*., 2011). A high-resolution cutoff was applied where CC½ (Karplus & Diederichs, 2012) in the highest-resolution shell reached approximately 0.3 (0.26–0.36 across datasets). This corresponded to resolution limits of 1.80 Å for the laser-off dark reference and the light-activated datasets (classes 4–7) and 1.70 Å for the merged pump–probe dark reference (classes 0–3); CC½ curves for all datasets are shown in Fig. S2. Final resolution cutoffs (1.85 Å for the laser-off and light-activated datasets and 1.70 Å for classes 0–3) were confirmed by monitoring Rfree during structure refinement. Dataset statistics are reported in Table 1.

**Table 1.** Data collection and refinement statistics.

| Data collection parameters | Dark state (laser-off) | Dark state (classes 0-3) | 0–350 $\mu\text{s}$ (175 $\mu\text{s}$) | 500–850 $\mu\text{s}$ (675 $\mu\text{s}$) | 1000–1350 $\mu\text{s}$ (1175 $\mu\text{s}$) | 1500–1850 $\mu\text{s}$ (1675 $\mu\text{s}$) |
| --- | --- | --- | --- | --- | --- | --- |
| Beamline | MicroMAX | MicroMAX | MicroMAX | MicroMAX | MicroMAX | MicroMAX |
| Photon energy | 13.7 keV | 13.7 keV | 13.7 keV | 13.7 keV | 13.7 keV | 13.7 keV |
| Temperature (K) | 293 | 293 | 293 | 293 | 293 | 293 |
| Detector | Jungfrau 9M | Jungfrau 9M | Jungfrau 9M | Jungfrau 9M | Jungfrau 9M | Jungfrau 9M |
| Monochromator | MLM | MLM | MLM | MLM | MLM | MLM |
| Relative energy bandwidth ($\Delta E/E$) | 1 % | 1 % | 1 % | 1 % | 1 % | 1 % |
| X-ray beam size (H x V) | $20 \times 5 \mu\text{m}^2$ | $20 \times 5 \mu\text{m}^2$ | $20 \times 5 \mu\text{m}^2$ | $20 \times 5 \mu\text{m}^2$ | $20 \times 5 \mu\text{m}^2$ | $20 \times 5 \mu\text{m}^2$ |
| Flux at sample position (photons $\text{s}^{-1}$) | $8.6 \times 10^{13}$ | $8.6 \times 10^{13}$ | $8.6 \times 10^{13}$ | $8.6 \times 10^{13}$ | $8.6 \times 10^{13}$ | $8.6 \times 10^{13}$ |
| X-ray pulse length | 350 $\mu\text{s}$ | 350 $\mu\text{s}$ | 350 $\mu\text{s}$ | 350 $\mu\text{s}$ | 350 $\mu\text{s}$ | 350 $\mu\text{s}$ |
| Repetition rate (X-ray/Laser) | 2 kHz/– | 2 kHz/– | 2 kHz/50 Hz | 2 kHz/50 Hz | 2 kHz/50 Hz | 2 kHz/50 Hz |
| Laser wavelength |  |  | 450 nm | 450 nm | 450 nm | 450 nm |
| Laser pulse energy | | | 3.3 $\mu\text{J}$ | 3.3 $\mu\text{J}$ | 3.3 $\mu\text{J}$ | 3.3 $\mu\text{J}$ |
| Laser pulse duration | | | 66 $\mu\text{s}$ | 66 $\mu\text{s}$ | 66 $\mu\text{s}$ | 66 $\mu\text{s}$ |
| Laser fluence at sample | – | | 61 $\text{mJ cm}^{-2}$ | 61 $\text{mJ cm}^{-2}$ | 61 $\text{mJ cm}^{-2}$ | 61 $\text{mJ cm}^{-2}$ |
| Nozzle size | 75 $\mu\text{m}$ | 75 $\mu\text{m}$ | 75 $\mu\text{m}$ | 75 $\mu\text{m}$ | 75 $\mu\text{m}$ | 75 $\mu\text{m}$ |
| Jet speed | 20 $\text{mm s}^{-1}$ | 20 $\text{mm s}^{-1}$ | 20 $\text{mm s}^{-1}$ | 20 $\text{mm s}^{-1}$ | 20 $\text{mm s}^{-1}$ | 20 $\text{mm s}^{-1}$ |
| Crystal size ($\mu\text{m}$) | ~15 $\mu\text{m}$ | ~15 $\mu\text{m}$ | ~15 $\mu\text{m}$ | ~15 $\mu\text{m}$ | ~15 $\mu\text{m}$ | ~15 $\mu\text{m}$ |
| **Data reduction** |  |  |  |  |  |  |
| Space group | P6<sub>5</sub>22 |  |  |  |  |  |
| Unit cell<br>*a, b, c*<br>$\alpha, \beta, \gamma$ | 122.57,<br>122.57,<br>46.60,<br>90, 90, 120 | 122.57,<br>122.57,<br>46.60,<br>90, 90, 120 | 122.57,<br>122.57,<br>46.60,<br>90, 90, 120 | 122.57,<br>122.57,<br>46.60,<br>90, 90, 120 | 122.57,<br>122.57,<br>46.60,<br>90, 90, 120 | 122.57, 122.57,<br>46.60,<br>90, 90, 120 |
| Resolution range | 46.61-1.80<br>(1.83-1.80) | 46.61-1.70<br>(1.73-1.70) | 46.61-1.80<br>(1.83-1.80) | 46.61-1.80<br>(1.83-1.80) | 46.61-1.80<br>(1.83-1.80) | 46.61-1.80<br>(1.83-1.80) |
| Total patterns | 767969 | 1080000 | 270000 | 270000 | 270000 | 270000 |
| Hits | 188544 | 880890 | 218251 | 218054 | 218057 | 217875 |
| Indexed patterns | 179320 | 611177 | 153964 | 154298 | 154303 | 154132 |
| Hit rate (%) | 24.55 | 81.56 | 80.83 | 80.76 | 80.76 | 80.69 |
| Indexing rate (%) | 95.11 | 69.38 | 70.54 | 70.76 | 70.76 | 70.74 |
| Completeness (%) | 100 (100) | 100 (100) | 100 (100) | 100 (100) | 100 (100) | 100 (100) |
| Redundancy | 6773.2<br>(4808.3) | 25136.9<br>(16987.4) | 6473.1<br>(4617.5) | 6465.0<br>(4609.8) | 6433.8<br>(4587.6) | 6424.4 (4583.0) |
| $R_{\text{split}}$ (%) | 3.58<br>(147.75) | 1.73<br>(137.8) | 3.06<br>(131.06) | 3.06<br>(131.66) | 3.18<br>(118.82) | 3.19 (128.18) |
| SNR | 16.13<br>(0.43) | 30.07<br>(0.52) | 17.97<br>(0.66) | 18 (0.62) | 17.94<br>(0.67) | 17.88 (0.63) |
| CC* | 1.000<br>(0.646) | 1.000<br>(0.658) | 1.000<br>(0.682) | 1.000<br>(0.722) | 1.000<br>(0.730) | 1.000 (0.709) |
| CC<sub>1/2</sub> | 0.998<br>(0.264) | 1.000<br>(0.276) | 0.999<br>(0.303) | 0.999<br>(0.352) | 0.999<br>(0.363) | 0.999 (0.335) |
| $R_{\text{iso}}$ (%) | – | – | 5.58 | 6.79 | 6.37 | 5.98 |
| **Refinement statistics** |  |  |  |  |  |  |
| Estimated active state contribution | N/A | N/A | 18 | 22 | 21 | 27 |
| Refinement resolution ($\text{\AA}$) | 42.67–1.85<br>(1.97-1.85) | 42.67–1.70<br>(1.78-1.70) | 42.67–1.85<br>(1.97-1.85) | 42.67–1.85<br>(1.97-1.85) | 42.67–1.85<br>(1.97-1.85) | 42.67–1.85<br>(1.97-1.85) |
| No. reflections | 18015 | 22626 | 16736 | 17132 | 16996 | 17445 |
| $R_{\text{Work}}$ | 0.171<br>(0.305) | 0.164<br>(0.43) | 0.214<br>(0.564) | 0.209<br>(0.546) | 0.211<br>(0.552) | 0.200 (0.519) |
| $R_{\text{Free}}$ | 0.187<br>(0.344) | 0.181<br>(0.416) | 0.235<br>(0.583) | 0.234<br>(0.562) | 0.238<br>(0.596) | 0.219 (0.501) |
| No. atoms |  |  |  |  |  |  |
| Total | 1012 | 972 | 982 | 966 | 956 | 950 |
| Protein | 892 | 892 | 887 | 878 | 874 | 874 |
| Water | 50 | 49 | 64 | 57 | 51 | 45 |
| Ligand/cofactor | 31 | 31 | 31 | 31 | 31 | 31 |
| Average atomic *B*-factor (Å<sup>2</sup>) |  |  |  |  |  |  |
| Overall | 50.0 | 49.48 | 42.7 | 43.5 | 42.85 | 44.98 |
| Protein | 49.6 | 48.86 | 42.2 | 43.2 | 42.57 | 44.77 |
| Water | 60.4 | 61.90 | 53.1 | 53.6 | 52.39 | 54.00 |
| Ligand/cofactor | 43.5 | 42.51 | 34.8 | 34.8 | 34.89 | 38.68 |
| Rmsd bond/ Rmsd angles | 0.007/0.94 | 0.006/0.857 | 0.01/1.16 | 0.01/1.14 | 0.008/1.126 | 0.007/1.112 |
| Ramachandran favoured/allowed/outliers | 99.07/0.93/0 | 100/0/0 | 99.06/0.94/0 | 100/0/0 | 100/0/0 | 100/0/0 |
| PDB code | 35GH | 35GI | 35GJ | 35GK | 35GL | 35GM |

### 2.6. Difference Fourier electron density maps

Fourier difference electron density maps were calculated using the *phenix.fobs_minus_fobs_map* program from the Phenix suite (Liebschner *et al*., 2019) with the multiscale scaling option enabled. Phases were derived from the refined laser-off dark-state model. Difference maps were generated by subtracting the merged pump–probe dark reference dataset (classes 0–3) from each light dataset corresponding to the four pump–probe delay times (classes 4–7).

To assess the presence of unintended photoactivation during pump–probe data collection, an additional difference Fourier map was calculated between the independently collected laser-off dark-state dataset and the merged pump–probe dark reference dataset (classes 0–3). The absence of significant difference density in the chromophore-binding pocket (Fig. 2*c*) was used to verify that the pump–probe dark reference was not affected by measurable light contamination and was therefore suitable for subsequent time-resolved analyses.

**Figure 2.**
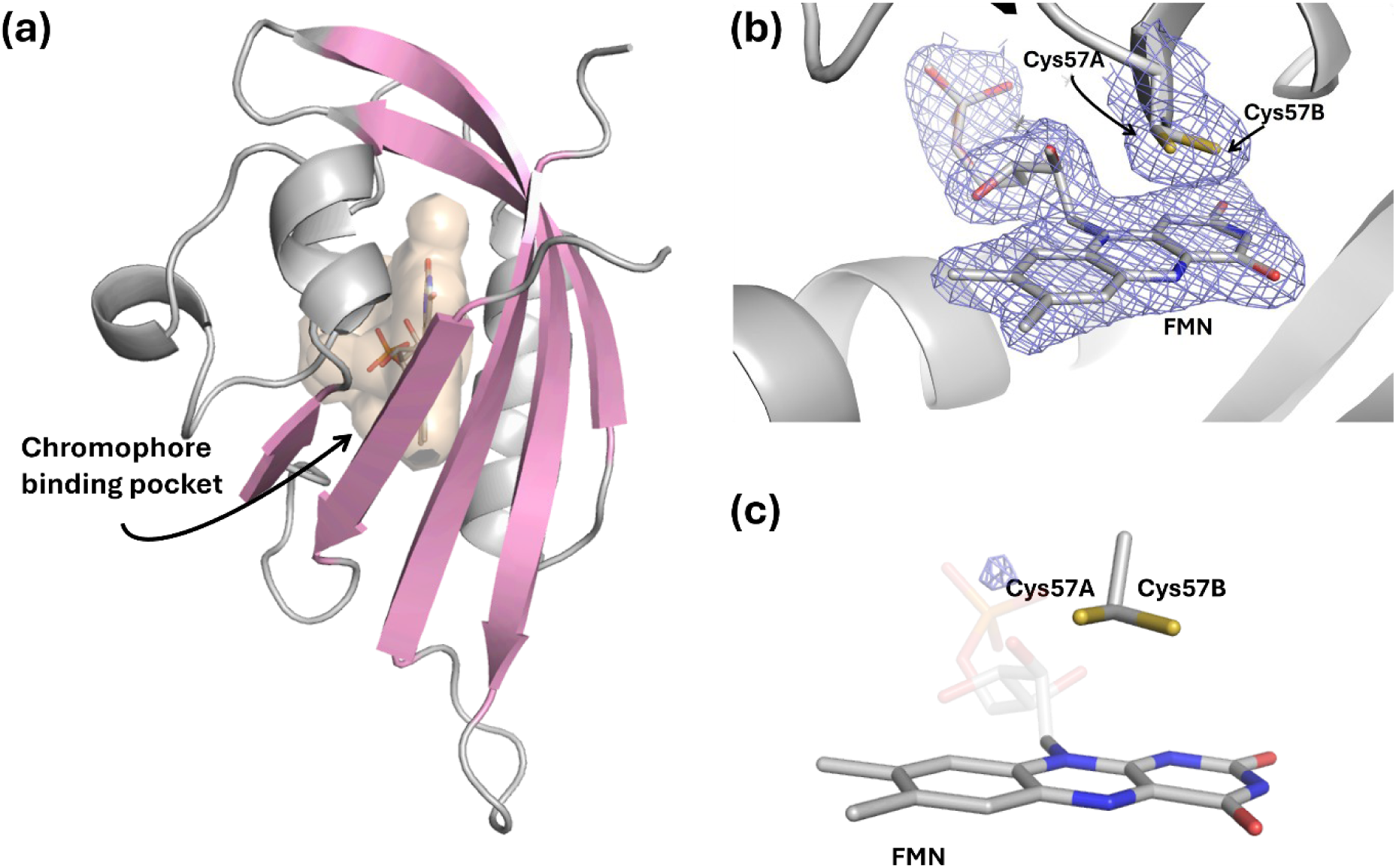
Dark-state structure of CrPhotLOV1 determined from the room-temperature laser-off dataset (1.85 Å). (*a*) Overall fold: the five antiparallel β-strands of the LOV core (pink) surround the FMN chromophore-binding pocket (FMN as sticks, pocket as surface). (*b*) 2*F*_o_ − *F*_c_ electron density (blue mesh) contoured at 1.5σ around FMN and Cys57, which is modeled in two conformations (Cys57A and Cys57B). (*c*) *F*_o_ − *F*_o_ difference map calculated by subtracting the laser-off structure factors from those of the pump–probe dark reference (classes 0–3), contoured at ±5σ using the same sign convention as the light-minus-dark maps in Fig. 3 (positive density in blue). No difference density is present at the Cys57 Sγ–FMN C4a adduct-forming site; a minor positive feature (∼6σ) lies near the terminal phosphate group of FMN.

### 2.7. Extrapolated electron density maps

The extrapolated structure factor amplitudes were calculated using a linear approximation (Genick *et al*., 1997) as follows: *F*_ext_ = [(*F*_obs_^light^ − *F*_obs_^dark^)/activated fraction] + *F*_obs_^dark^. Prior to extrapolation, the dark-state (classes 0–3) MTZ was truncated to 1.85 Å to match the refinement resolution of the light-state datasets. The same set of reflections was used as Rfree set for the dark-state and all light-state extrapolated datasets. An initial activated fraction was estimated by visual inspection of extrapolated electron density maps computed with phases from the refined dark-state model, selecting the highest extrapolation factor before the reappearance of density features corresponding to the dark-state conformation of Gln120, as in our previous work (Gotthard, Mous *et al*., 2024). The resulting 2*F*_ext_ − *F*_calc_ electron density maps showed features consistent with the corresponding *F*_obs_^light^ − *F*_obs_^dark^ difference Fourier maps, confirming the validity of the extrapolation. The corresponding light-state structure was then determined by rigid-body refinement against the extrapolated data, followed by manual removal of the dark-state Cys57 conformation in Coot and real-space and reciprocal-space refinement with covalent restraints for the Cys57–FMN C4a photoadduct and modified FMN planarity. The activated fraction was validated *a posteriori* using the refined light-state model. Zero-cycle refinement was performed against each extrapolated MTZ to obtain properly weighted difference map coefficients, and the F_o_-F_c_ density at the Cys57 Sγ adduct position was evaluated as a function of α using an in-house shell-based workflow. The upper bound on α was established as the value at which this density changed sign from positive to negative. The activated fractions used for the final extrapolation were α = 0.18, 0.22, 0.21 and 0.27 for classes 4, 5, 6 and 7, respectively, with upper bounds of α ≈ 0.19, 0.23, 0.23 and 0.29 determined by the adduct density sign-change criterion (Fig. S3).

In parallel, extrapolated structure factor amplitudes and electron density maps were also generated using the program Xtrapol8 (De Zitter *et al*., 2022). Xtrapol8 performs automated extrapolation of structure factor amplitudes and facilitates the generation of extrapolated maps and refinement of light-state models using occupancy-dependent extrapolation procedures. The resulting extrapolated maps and refined models were compared with those obtained using the linear extrapolation approach to evaluate whether both extrapolation strategies converged on the same structural interpretation of the light-activated state.

### 2.8. Model building and refinement

The dark-state structure was determined by molecular replacement using Phaser (McCoy *et al*., 2007) with the LOV1 domain of Chlamydomonas reinhardtii (PDB: 1N9L; Fedorov *et al*., 2003) as a search model. The resulting solution was refined against the laser-off dark reference dataset to 1.85 Å resolution through iterative cycles of reciprocal-space refinement in PHENIX (Liebschner *et al*., 2019) and manual model building in Coot (Emsley *et al*., 2010), including optimization of side-chain conformations and placement of water molecules. The refined laser-off dark-state model was subsequently used as the starting model for rigid-body refinement in PHENIX against the merged pump–probe dark reference dataset (classes 0–3) to 1.70 Å resolution, yielding an independent dark-state structure for the pump–probe experiment. Light-state structures in each pump–probe time window (0–350, 500–850, 1000–1350 and 1500–1850 µs) were refined to 1.85 Å resolution against extrapolated structure factor amplitudes derived from the corresponding light-activated datasets (classes 4–7), as described above. Rigid-body refinement was performed first, followed by restrained reciprocal-space and real-space refinement with occupancies of the light-state conformer refined freely. Model representation and analysis were prepared with PyMOL (https://pymol.org/).

## 3. Results

### 3.1. Overall data quality and dataset consistency

High-quality room-temperature serial synchrotron crystallography datasets were collected for an independently measured laser-off dark-state reference dataset and for the pump–probe experiment, comprising the merged dark reference (classes 0–3) and four light-activated time windows (0–350, 500–850, 1000–1350 and 1500–1850 µs). All datasets exhibited consistent crystallographic statistics, including high completeness, redundancy, and comparable resolution limits (Table 1). No measurable differences in unit-cell parameters were observed among the independent dark-state dataset and all pump–probe datasets, with unit-cell dimensions remaining indistinguishable within experimental uncertainty. This excellent isomorphism provided the basis for reliable difference Fourier analysis and calculation of extrapolated structure factor amplitudes, enabling direct structural comparison between the independently refined dark-state model and the light-activated pump–probe datasets.

Stable crystal delivery with the injector-based pump–probe setup (Fig. 1) resulted in reproducible data collection across all time points. Inspection of the dark-state electron density maps revealed no evidence of site-specific damage at the FMN chromophore or Cys57, both of which are known to be sensitive to radiation induced changes. This indicates that the X-ray dose used in the experiments did not cause detectable damage at these locations. Together, these results demonstrate the robustness and reproducibility of the injector-based workflow implemented at the MicroMAX beamline.

### 3.2. Dark-state structure and verification of the absence of light contamination

The high quality of the dark-state dataset enabled unambiguous modeling of the FMN chromophore and the surrounding active-site residues. The 2Fo–Fc electron density clearly defines the chromophore-binding pocket, with Cys57 modeled in two conformations that are fully supported by the experimental density (Fig. 2).

A critical aspect of injector-based time-resolved serial crystallography is ensuring that the crystals remain in the dark state until photoexcitation, i.e. that the dark reference dataset used for Fo–Fo difference-map calculations and extrapolated structure factor analysis remains free from unintended photoactivation. To verify this, the merged pump–probe dark reference dataset (classes 0–3), corresponding to diffraction images collected prior to laser excitation, was compared with an independently collected laser-off dataset acquired without optical illumination. Difference Fourier maps calculated between these datasets were inspected for residual signal in the chromophore-binding pocket. No positive or negative difference density was detected at the Cys57 Sγ–FMN C4a position in the dark-reference map (Fig. 2*c*), whereas the light-state maps showed a pronounced positive feature at this site, reaching ∼21 σ at Cys57 Sγ and ∼18 σ at FMN C4a (Fig. 3*e*). Although a residual positive peak of ∼6 σ was present near the terminal phosphate group of FMN, it was spatially distinct from the adduct-forming region. These results demonstrate that the pump–probe dark reference is consistent with the true dark state, with no detectable evidence of light contamination at the FMN binding site.

**Figure 3.**
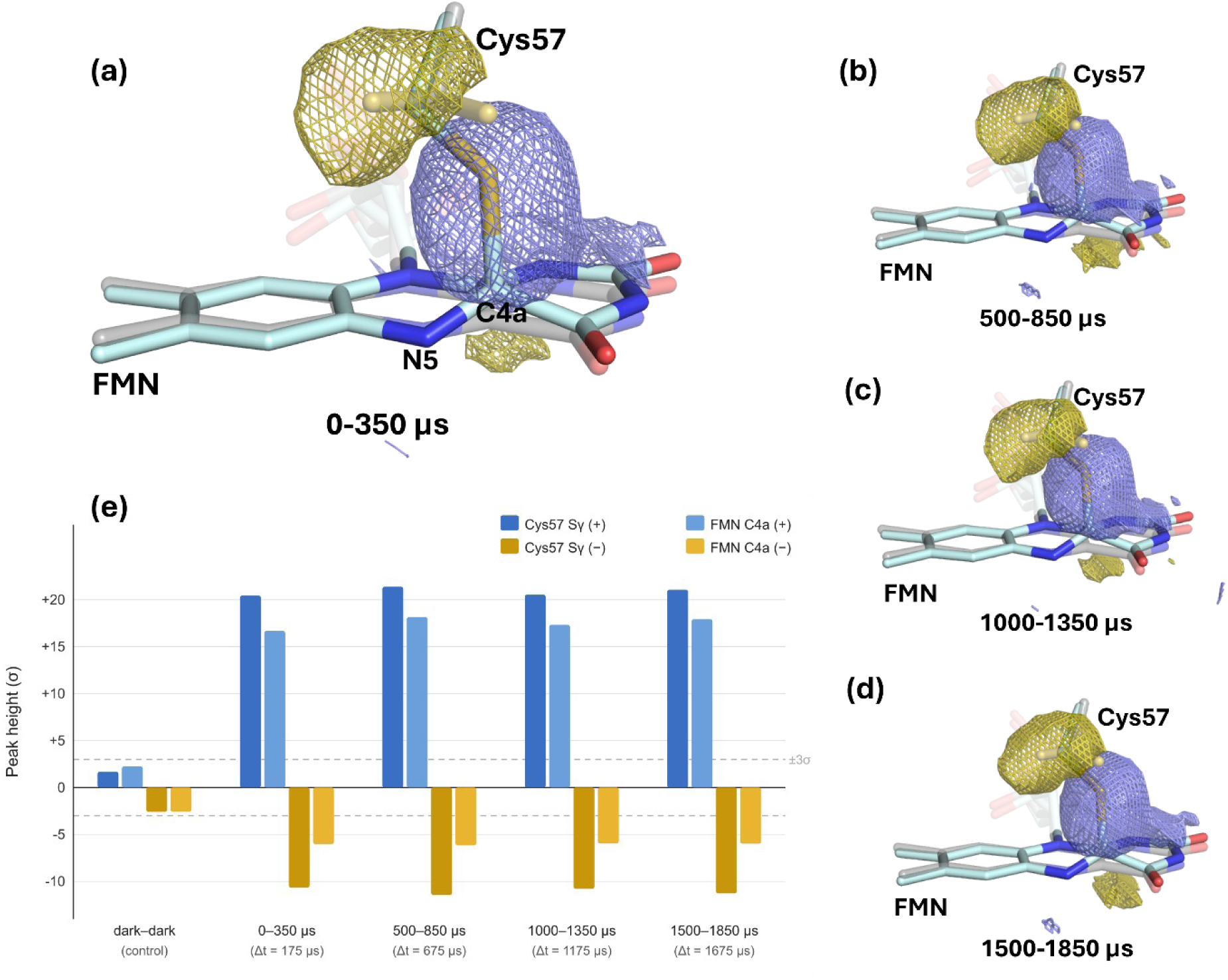
Photoadduct formation in CrPhotLOV1 revealed by *F*_o_ − *F*_o_ difference maps. (*a*)–(*d*) Light-minus-dark maps (classes 4–7 minus the pump–probe dark reference, classes 0–3) for the time windows (*a*) 0–350 µs, (*b*) 500–850 µs, (*c*) 1000–1350 µs and (*d*) 1500–1850 µs, contoured at ±5σ (positive density in blue, negative in yellow). Phases for all difference maps were derived from the refined laser-off dark-state model. The refined light-state models for the respective windows (cyan sticks) are overlaid on the dark-state model (grey). (*e*) Peak heights of the positive (+) and negative (−) difference density at the Cys57 Sγ and FMN C4a positions in the maps in (*a*)–(*d*) and in the control map between the laser-off dataset and the pump–probe dark reference (dark–dark; Fig. 2c). Dashed lines mark ±3σ. In the control map all peaks are within ±2.6σ; in the four light-minus-dark maps the positive peaks exceed 16σ at FMN C4a and 20σ at Cys57 Sγ.

Comparison of the dark-state model with previously reported CrPhotLOV1 structure revealed close agreement in both the overall fold and the architecture of the chromophore-binding pocket, with an RMSD of 0.15 Å relative to the dark-state structure from our previous study (Gotthard, Mous *et al*., 2024; PDB 8QI9).

### 3.3. Validation of photoactivation by difference Fourier analysis

Successful light activation during pump–probe data collection was verified using Fo–Fo difference Fourier maps calculated relative to the pump–probe dark reference (classes 0–3) (Figs. 3*a*–3*d*).

Following laser excitation, clear difference density appeared around the FMN chromophore and the photoactive residue Cys57 at all investigated delay times. Fourier difference electron density maps calculated for the four pump–probe time windows (0–350, 500–850, 1000–1350 and 1500–1850 µs) revealed reproducible positive and negative features within the chromophore-binding pocket, demonstrating successful photoactivation under the applied experimental conditions (Figs. 3*a*–3*d*). The overall pattern of difference density was consistent across all time points, with the most prominent signals localized around the FMN chromophore and the photoactive residue Cys57. A strong positive peak positioned between the sulfur atom of Cys57 and the C4a atom of FMN, together with a corresponding negative peak at the distal dark-state position of the cysteine side chain, is consistent with formation of the characteristic cysteinyl–FMN thioether adduct. In the dark state, Cys57 is present in two alternative conformations, and photoactivation is accompanied by a reorientation of its side chain about the Cα–Cβ bond. The close similarity of the four maps is expected from the photocycle kinetics of CrPhotLOV1: adduct formation is complete within about 20 µs of excitation and the adduct persists for hundreds of seconds (Kottke *et al*., 2003), so all four delays probe the same photoadduct state and the four datasets represent independent replicate measurements of this state.

Additional difference density features were observed throughout the FMN molecule, reflecting local structural rearrangements accompanying adduct formation. Positive and negative peaks above and below the isoalloxazine ring are consistent with distortion of the flavin geometry following covalent attachment, while pronounced difference density associated with the phosphate moiety indicates a local repositioning of the chromophore that preserves its overall anchoring within the binding pocket. Significant positive and negative difference density was also observed for the conserved residue Gln120, consistent with reorientation of its side chain and reorganization of the hydrogen-bonding network surrounding the chromophore following photoactivation.

### 3.4. Extrapolated electron density maps and light-state structural models

To facilitate structural interpretation of the light-induced structural changes, extrapolated structure factor amplitudes were calculated for each pump–probe time point and used to generate extrapolated electron density maps. The resulting maps were of sufficient quality to enable model building and refinement of light-state structures (Fig. 4*a*, Fig. S4).

**Figure 4.**
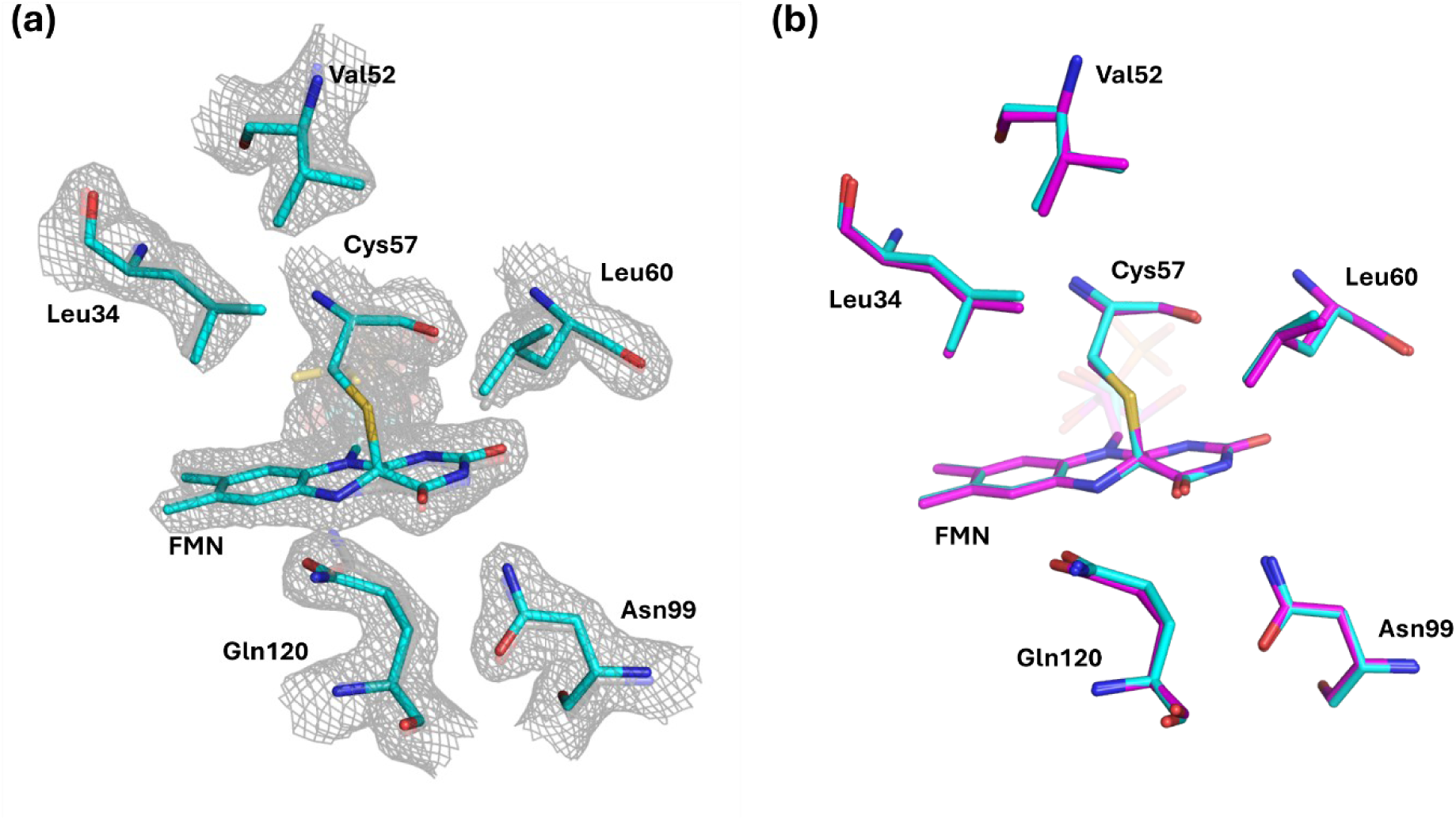
Structural changes on photoactivation of CrPhotLOV1. (*a*) Extrapolated electron density (2*F*_ext_ − *F*_c_ map, α = 0.27; grey mesh) contoured at 1.5σ for the 1500– 1850 µs time window, with the refined dark-state (grey sticks) and light-state (cyan) models. The map supports the covalent FMN–Cys57 thioether adduct and local changes, including rotation of the Gln120 side chain and small shifts of Leu34, Val52, Leu60 and Asn99. (*b*) Superposition of the light-state model from this study (cyan; 1500–1850 µs) with the millisecond light-state structure of CrPhotLOV1 (magenta; 0–5 ms) (Gotthard, Mous *et al*., 2024; PDB ID: 8QIA). The two models agree closely in the chromophore-binding pocket, including the FMN–Cys57 adduct and the rotated Gln120 side chain.

Structural refinement against the extrapolated structure factor amplitudes confirmed the interpretation of the difference Fourier maps. Well-defined electron density supporting formation of the covalent FMN–Cys57 adduct was obtained even at relatively low extrapolation factors, demonstrating that extrapolation enhances a genuine light-induced structural signal rather than introducing modelling artefacts.

Refinement of extrapolated light-state models demonstrated that the collected datasets support reliable structural interpretation of the photoactivated state. The refined models revealed localized conformational rearrangements within the chromophore-binding pocket (Fig. 4*a*). The main structural changes are localized around the FMN chromophore and the photoactive residue Cys57, consistent with photoadduct formation and the accompanying reorganization of the local hydrogen-bonding network. In particular, the conserved residue Gln120 adopts a rotated amide orientation relative to the dark state, repositioning its side chain toward the protonated N5 atom of the FMN chromophore. This rearrangement is fully supported by the extrapolated electron density and is consistent with the prominent positive and negative features observed in the corresponding difference Fourier maps. In addition to the chromophore and Gln120, local conformational adjustments were required for several neighbouring residues, most notably Leu34 and Val52, whose refined side-chain conformations follow the experimentally observed difference density. The locations of these rearrangements coincide with the strongest peaks in the Fo–Fo maps, providing independent support for the refined light-state models. In contrast, the overall protein architecture remains essentially unchanged throughout the microsecond time range investigated, indicating that the observed response is largely confined to the immediate environment of the chromophore.

Together, the difference Fourier maps and the extrapolated structural models demonstrate that the MicroMAX pump–probe workflow enables reliable time-resolved structural characterization of light-induced structural changes in CrPhotLOV1.

### 3.5. Comparison of extrapolation methods

To test whether the structural interpretation depends on the estimated activation levels, extrapolation was repeated with Xtrapol8. Extrapolated electron density maps generated using Xtrapol8, with a uniform activation level of α = 0.18 across all four time windows, yielded refined light-state models fully consistent with those obtained using the per-class activation levels determined by the sign-change criterion (α = 0.18, 0.22, 0.21 and 0.27 for classes 4–7, respectively). Both approaches reproduced the characteristic FMN–Cys57 covalent adduct, rotation of the conserved Gln120 side chain, and local conformational rearrangements involving Leu34 and Val52, while preserving the overall protein architecture. For the three earlier time windows (classes 4–6), the two activation estimates agree to within 0.04, and the resulting maps are essentially indistinguishable. For the latest time window (class 7), the larger discrepancy (Δα = 0.09) produces modestly noisier extrapolated maps with the Xtrapol8 value, yet the structural interpretation remains unchanged. The subsequent analysis is therefore presented using the per-class activation levels for consistency throughout the study.

### 3.6. Assessment of photoactivation levels

The activation levels estimated for the four pump–probe datasets ranged from 18% to 27% (Table 1), indicating comparable levels of photoactivation across all investigated time points. Although these values are lower than the maximum activation level reported for the HVE-based experiment at SLS (65%) (Gotthard, Mous *et al*., 2024), they fall well within the range observed across the full time series (15-65%). Despite minor variations between datasets, all activation levels were sufficient to generate high-quality extrapolated structure factor amplitudes, enabling reliable map calculation, model refinement, and structural interpretation of the light-activated state. The comparable activation levels obtained throughout the microsecond time range demonstrate stable and reproducible optical excitation under the applied experimental conditions, supporting direct comparison of the refined light-state structures. Together, these results further demonstrate the robustness of the injector-based pump–probe workflow implemented at the MicroMAX beamline.

### 3.7. Comparison with previously reported light-state structures

The refined light-state structures closely resemble previously reported light-activated CrPhotLOV1 models determined at millisecond time delays (Gotthard, Mous *et al*., 2024; PDB 8QIA). In particular, formation of the characteristic FMN–Cys57 thioether adduct and rotation of the conserved Gln120 side chain are reproduced, together with the overall organization of the chromophore-binding pocket (Fig. 4*b*). The agreement between independently determined structures provides strong support for the validity of the present experimental workflow and structural interpretation.

## 4. Discussion

The present study demonstrates the successful implementation of an injector-based pump–probe serial synchrotron crystallography workflow at the MicroMAX beamline for time-resolved structural studies of the LOV photoreceptor CrPhotLOV1. The high quality and excellent isomorphism of the serial datasets enabled reliable calculation of both difference Fourier maps and extrapolated electron density maps, providing a robust basis for structural interpretation of light-induced conformational changes. Importantly, independent dark-state measurements confirmed the absence of detectable light contamination during pump–probe data collection and the integrity of the reference dataset.

The observed structural changes are fully consistent with the expected LOV-domain photochemistry. Difference Fourier maps and refined extrapolated models consistently revealed formation of the characteristic cysteinyl–FMN thioether adduct, together with localized rearrangements involving the conserved residue Gln120 and neighbouring side chains. These structural changes remained confined to the chromophore-binding pocket, while the overall protein architecture was essentially unchanged throughout the investigated microsecond time range. Comparable structural models obtained using both the linear extrapolation procedure and Xtrapol8 further demonstrate that the structural interpretation is robust with respect to the extrapolation strategy, despite differences in the estimated activation levels.

Several aspects warrant further discussion. First, the four pump–probe datasets are expected to represent the same long-lived photoactivated state: in CrPhotLOV1 the FMN triplet state decays into the thioether adduct with time constants of 0.8 and 4 µs and the adduct persists for hundreds of seconds (Kottke *et al*., 2003). Therefore, no time-dependent structural changes are expected within the 0–1850 µs range. Rather, the four independently collected datasets demonstrate the reproducibility of activation levels, difference density and refined structural models. Second, activation levels of 18 to 27% are moderate. Contributing factors likely include the Gaussian laser profile, which has an 80 µm diameter comparable to that of the extruded stream so that crystals at the stream periphery receive only a fraction of the peak fluence, light scattering in the viscous matrix (Niziński *et al*., 2025), and the high optical density of flavin-containing crystals at 450 nm, which limits light penetration into 10–20 µm crystals. Because these factors are expected to be similar in other HVE-based experiments, differences in excitation pulse duration and energy may also contribute to the lower activation compared with the earlier experiment at SLS. A systematic variation of the pulse energy, together with an estimate of the photon dose per chromophore, would be needed to establish that excitation was in the linear regime and to exclude multiphoton effects that have been documented for high-fluence femtosecond pump–probe experiments at XFELs (Grünbein *et al*., 2020; Barends *et al*., 2024; Bertrand *et al*., 2024). Third, the frame-based dark–light scheme uses only a fraction of the recorded frames, because frames recorded while the illuminated sample volume still passes the X-ray interaction point cannot serve as dark reference; the stream velocity of approximately 20 µm/ms defines the time after which fresh sample is probed, and the corresponding sample consumption should be weighed against the simplicity of the setup. Finally, the present measurements at microsecond delays complement our previous TR-SSX study of CrPhotLOV1 at millisecond delays (Gotthard, Mous *et al*., 2024) and the TR-SFX series at XFELs (Olasz *et al*., 2026); together they show that injector-based TR-SSX with the same crystal preparation can be transferred between synchrotron beamlines with consistent structural results. With respect to time resolution, the 0–350 µs window can be compared with earlier injector-based measurements: the 250–750 µs snapshot of the xenorhodopsin BcXeR at P14 (PETRA III) used a 500 µs exposure and a 500 µs pump pulse that overlapped the probe window by 250 µs (Kovalev *et al*., 2023), following the 5 ms frames of the earlier bacteriorhodopsin study at SLS (Weinert *et al*., 2019). The microsecond excitation used here defines the reaction start to within the frame timing, and the 350 µs exposure is shorter and starts earlier than the 500 µs exposure of the BcXeR snapshot; the 66 µs pump overlaps only the first 66 µs of the 0–350 µs window, and a laser-off control is retained.

The combination of stable injector-based sample delivery, reproducible optical excitation, high-quality serial diffraction data and reliable extrapolation procedures establishes a practical workflow for routine time-resolved serial synchrotron crystallography experiments at MicroMAX. This protocol should be readily applicable to other photoactive proteins and provides a solid framework for future time-resolved studies of light-driven structural dynamics.

Since the experiments reported here, the light-triggered serial crystallography setup has been further developed at MicroMAX. The pump laser can now be delivered in a quasi-collinear geometry relative to the X-ray beam (Fig. S5), facilitating spatial overlap at the sample. Together with the tunable nanosecond laser system, modulated laser diodes provide flexible excitation from nanosecond to microsecond–millisecond timescales. New user tools, including the integration of the laser alignment into MXCuBE (Mueller *et al*., 2017), have been developed to enable visualization and verification of the laser position, adjustment of laser power and pulse duration, and switching between dark and time-resolved acquisition modes. In addition, KB mirrors are being commissioned to provide higher photon flux and a smaller X-ray focus (Gonzalez *et al*., 2026), potentially enabling shorter exposure times and extending TR-SSX at MicroMAX towards shorter timescales.

## Acknowledgements

We acknowledge the MAX IV Laboratory for beamtime on the MicroMAX beamline under proposal 20251858. Research conducted at MAX IV, a Swedish national user facility, is supported by Vetenskapsrådet (Swedish Research Council, VR) under contract 2018-07152, Vinnova (Swedish Governmental Agency for Innovation Systems) under contract 2018-04969 and Formas under contract 2019-02496. MicroMAX is funded by the Novo Nordisk Foundation under the grant number NNF17CC0030666. We acknowledge the P11 beamline of PETRA III operated by DESY Photon Science (No. R-20240820 EC) for cryo-crystallography beamtime carried out in preparation for this experiment. We acknowledge the European Synchrotron Radiation Facility (ESRF) for provision of synchrotron radiation facilities under proposals MX-2593 and MX-2692, and we would like to thank the beamline staff for assistance and support. We acknowledge the Malopolska Centre of Biotechnology Structural Biology Core Facility for access and support. We thank the Department of Microbiology, Faculty of Biochemistry, Biophysics and Biotechnology, Jagiellonian University, for access to its bacterial culture and cell disruption infrastructure. We thank Michal Kepa (Paul Scherrer Institute, Villigen, Switzerland) for his contribution to the laser-diode setup at MicroMAX.

The authors used Claude (Anthropic) and Microsoft Copilot for language editing and text refinement. AI-assisted tools were additionally used in the modification of a schematic figure. All scientific analyses, data interpretation, and final manuscript content were performed, reviewed, and approved by the authors.

## Funding information

The project was financed under Dioscuri, a program initiated by the Max Planck Society, jointly managed with the National Science Centre in Poland, and mutually funded by the Polish Ministry of Science and Higher Education and the German Federal Ministry of Research, Technology and Space. This research was funded by the Polish National Science Centre (grant agreement No. UMO-2021/03/H/NZ1/00002 and SONATA BIS grant No. UMO-2024/54/E/NZ1/00429, both awarded to P.N.). The purchase of equipment used in the project has been supported by a grant of the Faculty of Biochemistry, Biophysics and Biotechnology under the Strategic Programme Excellence Initiative at Jagiellonian University. The work presented in this paper benefits from the support received from the Polish Ministry of Science and Higher Education, dec. no. 2021/WK/11.

## Author contributions

P.N. conceived and coordinated the study. J.N. coordinated the beamline preparation and supported the experiment. B.O., A.K., A.S., M.P. and A.A. expressed, purified and crystallized CrPhotLOV1. A.K. and A.S. optimized crystal injection conditions. C.M.C., S.M.K., L.L.C., D.L., M.M., T.U., M.C. and J.N. prepared and operated the beamstation, including the laser systems. C.M.C., B.O. and M.K.S. performed data processing during and after beamtime. M.W. refined and interpreted the structures. M.W. and P.N. wrote the manuscript with direct contributions from J.N., M.C. and B.O., and further contributions from all co-authors. All authors read and approved the final manuscript.

## Conflict of interest

The authors declare no conflicts of interest.

## Data availability

Coordinates and structure factors have been deposited in the Protein Data Bank with the following accession codes: dark state, laser-off (35GH); dark state, classes 0–3 (35GI); and light states for the 0–350 µs (35GJ), 500–850 µs (35GK), 1000–1350 µs (35GL) and 1500–1850 µs (35GM) windows after photoexcitation.

---

**Figure S1.**
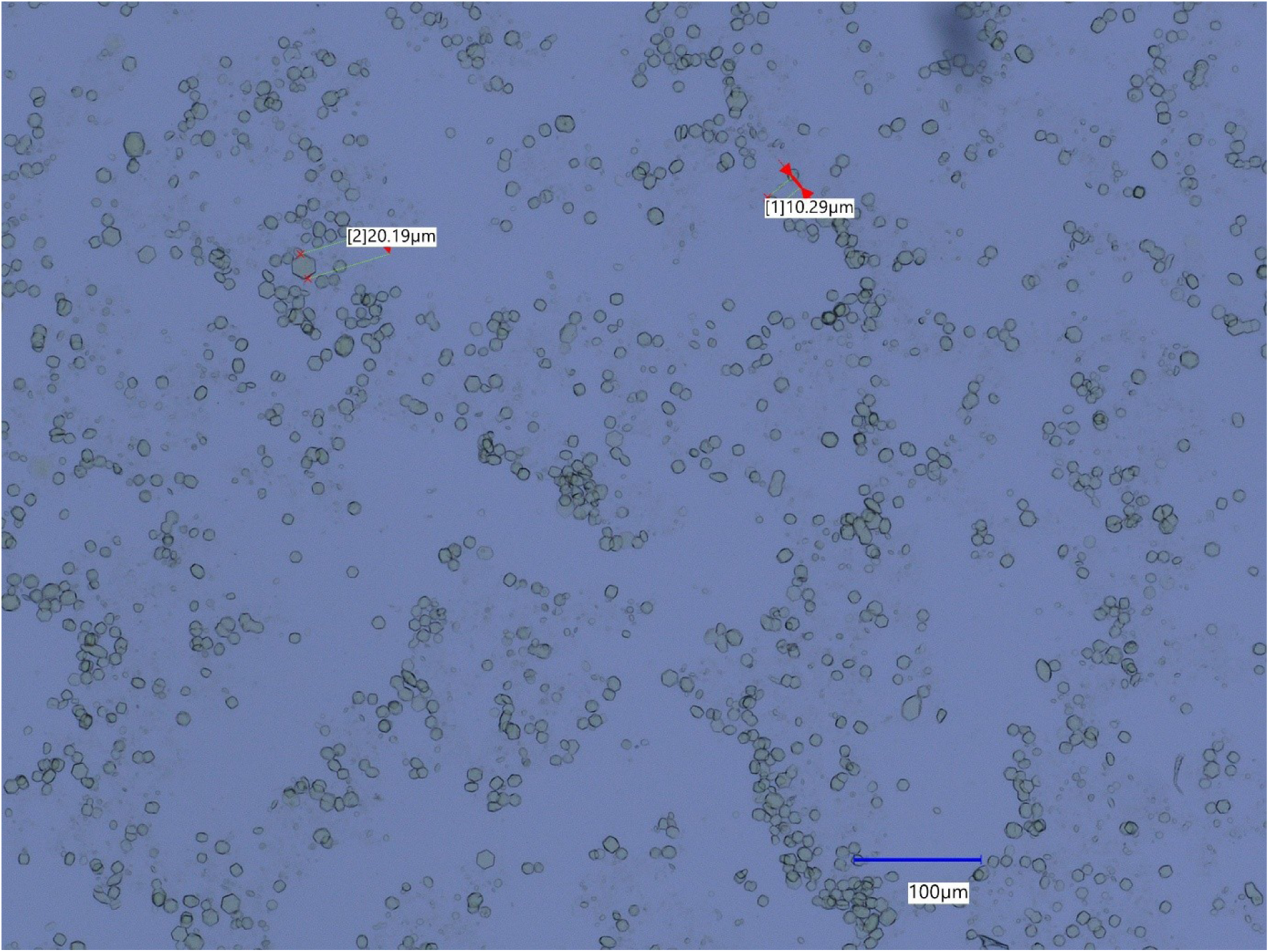
Micrograph of CrPhotLOV1 microcrystals. Crystals were imaged by light microscopy prior to data collection. Representative crystal sizes of 10–20 µm are indicated (red arrows). Scale bar: 100 µm.

**Figure S2.**
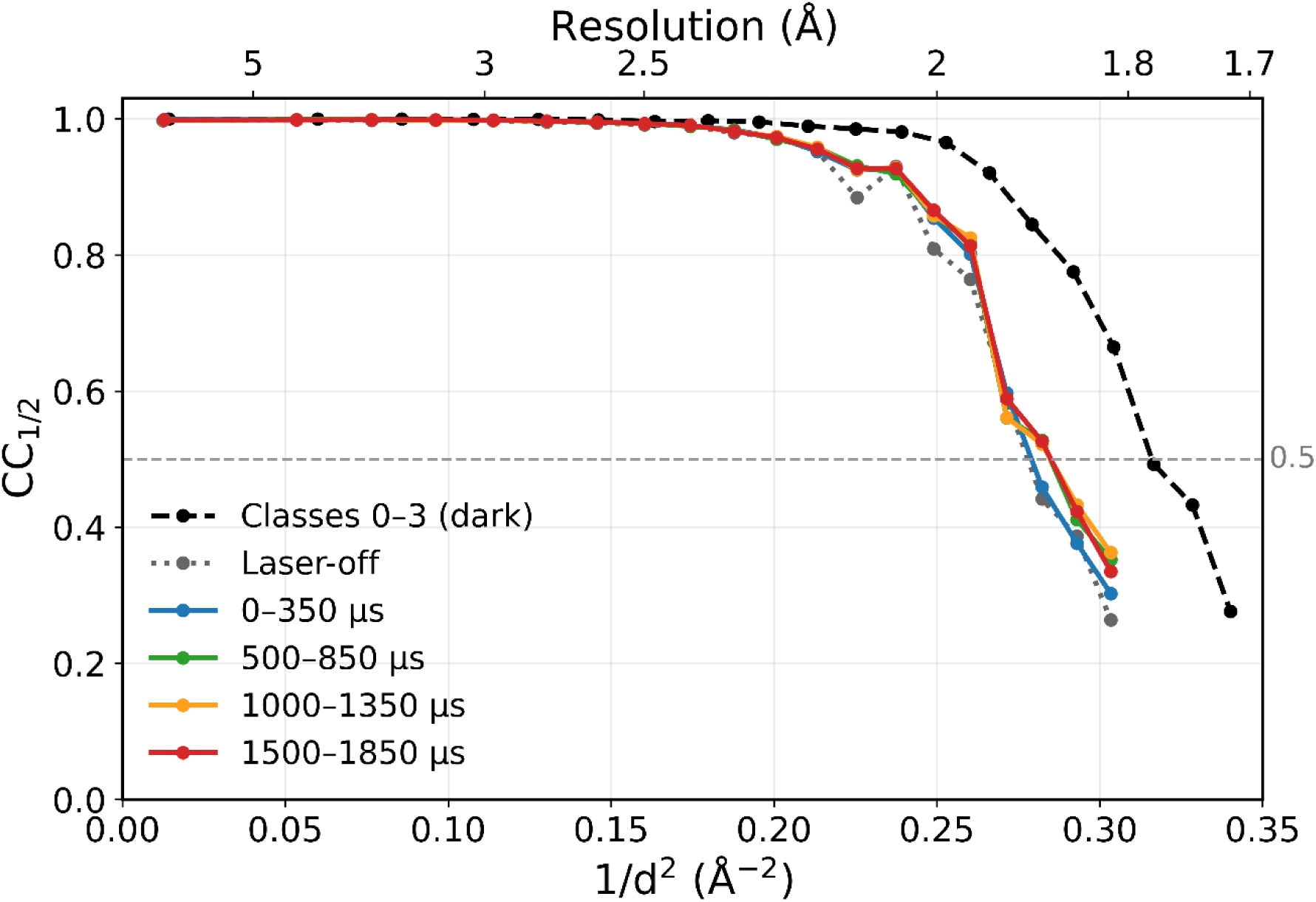
CC½ as a function of resolution (plotted against 1/*d*^2^) for the merged pump– probe dark reference, classes 0–3 (black, dashed), the laser-off dark reference (grey, dotted) and the light-activated classes for the time windows 0–350 µs (blue), 500–850 µs (green), 1000–1350 µs (orange) and 1500–1850 µs (red). The grey dashed horizontal line marks CC½ = 0.5. The high-resolution limits were set to 1.80 Å for the laser-off and light-activated datasets and 1.70 Å for classes 0–3, where CC½ in the highest-resolution shell was 0.26–0.36; see Section 2.5.

**Figure S3.**
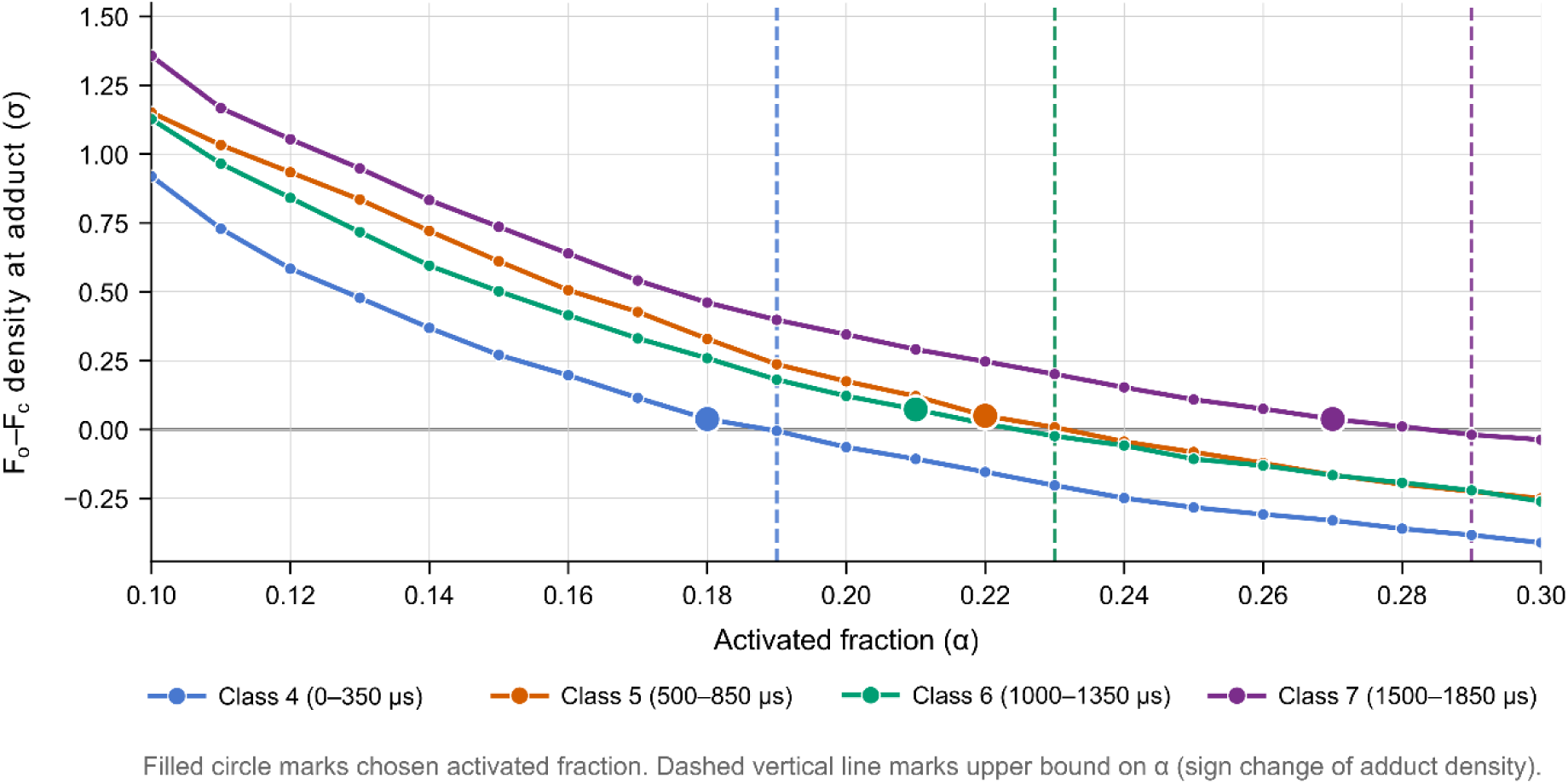
Activation-level scan for the light-activated classes 4–7. *F*_o_ − *F*_c_ density (σ) at the Cys57 Sγ adduct position as a function of the activated fraction α, from zero-cycle refinement of the light-state model against the extrapolated data. Class 4 (0–350 µs), blue; class 5 (500–850 µs), orange; class 6 (1000–1350 µs), green; class 7 (1500–1850 µs), purple. Filled circles mark the activated fraction used for the final extrapolation (α = 0.18, 0.22, 0.21 and 0.27, respectively); dashed vertical lines mark the upper bound on α, at which the adduct density changes sign from positive to negative. Classes 5 and 6 share the same sign-change value (α = 0.23); their dashed lines coincide.

**Figure S4.**
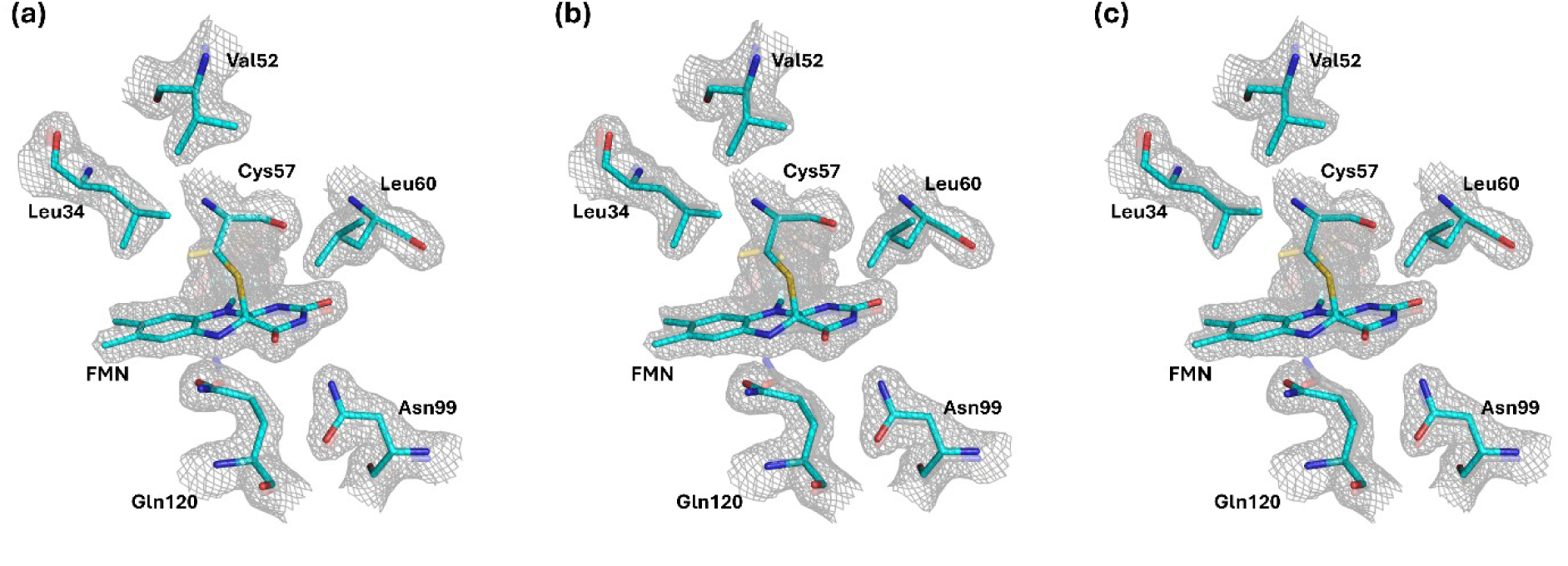
Extrapolated electron density (2*F*_ext_ − *F*_c_ map; grey mesh) contoured at 1.5σ for the (*a*) 0–350 µs (α = 0.18), (*b*) 500–850 µs (α = 0.22) and (*c*) 1000–1350 µs (α = 0.21) time windows, with the refined dark-state (grey sticks) and light-state (cyan sticks) models. The maps support the covalent FMN–Cys57 thioether adduct and local structural changes, including rotation of the Gln120 side chain and small shifts of Leu34, Val52, Leu60 and Asn99.

**Figure S5.**
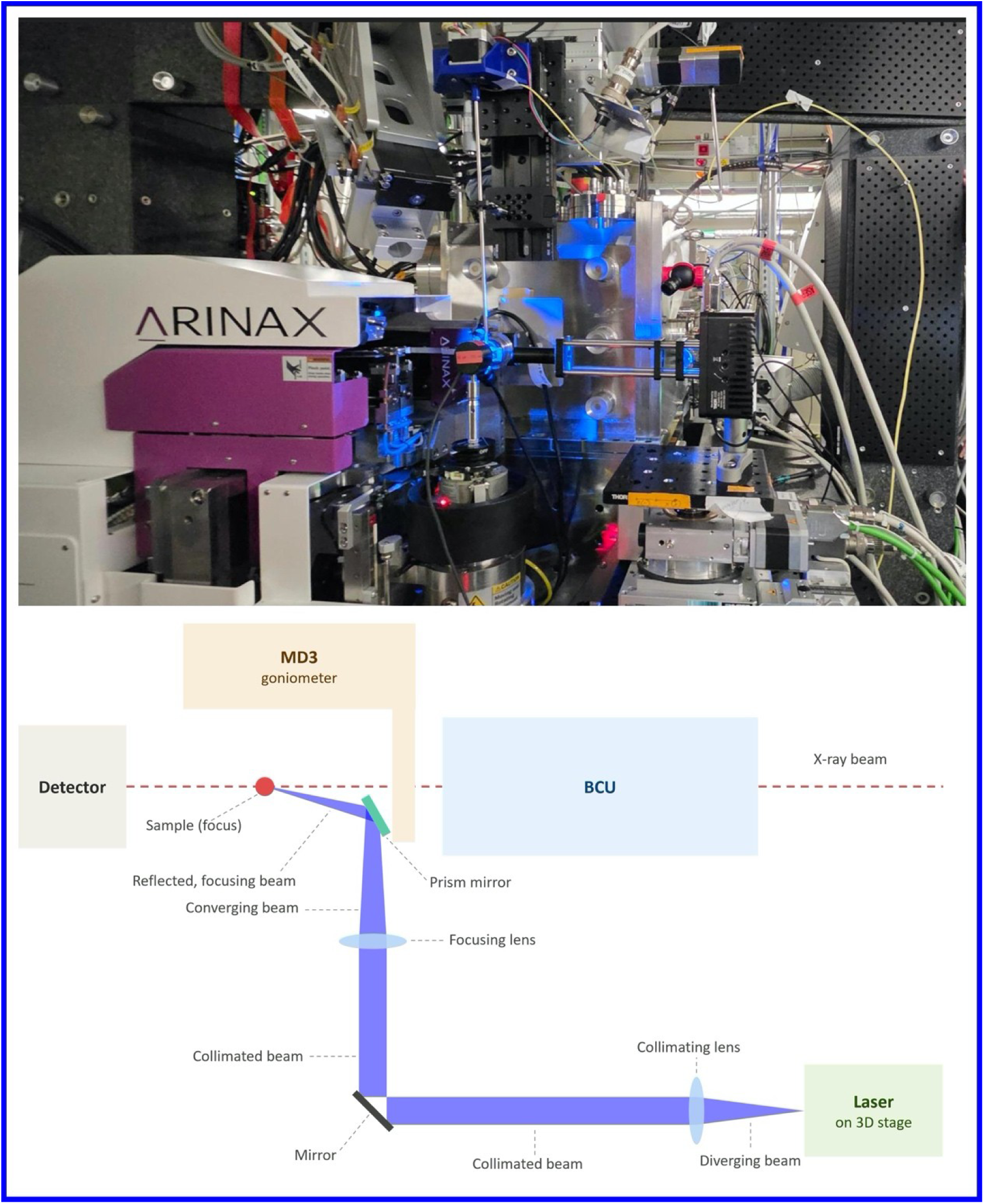
Quasi-collinear laser–X-ray geometry at MicroMAX. Top: Current laser delivery setup used for an HVE injector-based TR-SSX measurement. Bottom: Schematic top view of the laser delivery geometry and the main experimental components, including the beam conditioning unit (BCU), MD3 microdiffractometer, detector, and laser mounted on a 3D translation stage. The schematic illustrates the collimation, focusing, and reflection of the laser beam toward the sample interaction point.

## Notes

### Competing Interest Statement

The authors have declared no competing interest.

## References

Aumonier, S., Santoni, G., Gotthard, G., von Stetten, D., Leonard, G. A. & Royant, A. (2020). IUCrJ 7, 728–736.

Barends, T. R. M., Gorel, A., Bhattacharyya, S., Schirò, G., Bacellar, C., Cirelli, C., Colletier, J.-P., Foucar, L., Grünbein, M. L., Hartmann, E., Hilpert, M., Holton, J. M., Johnson, P. J. M., Kloos, M., Knopp, G., Marekha, B., Nass, K., Nass Kovacs, G., Ozerov, D., Stricker, M., Weik, M., Doak, R. B., Shoeman, R. L., Milne, C. J., Huix-Rotllant, M., Cammarata, M. & Schlichting, I. (2024). Nature 626, 905–911.

Bertrand, Q., Nogly, P., Nango, E., Kekilli, D., Khusainov, G., Furrer, A., James, D., Dworkowski, F., Skopintsev, P., Mous, S., Martiel, I., Börjesson, P., Ortolani, G., Huang, C.-Y., Kepa, M., Ozerov, D., Brünle, S., Panneels, V., Tanaka, T., Tanaka, R., Tono, K., Owada, S., Johnson, P. J. M., Nass, K., Knopp, G., Cirelli, C., Milne, C., Schertler, G., Iwata, S., Neutze, R., Weinert, T. & Standfuss, J. (2024). Nat. Commun. 15, 10278.

Bosman, R., Ortolani, G., Ghosh, S., James, D., Norder, P., Hammarin, G., Úlfarsdóttir, T. B., Ostojić, L., Weinert, T., Dworkowski, F., Tomizaki, T., Standfuss, J., Brändén, G. & Neutze, R. (2025). Nat. Commun. 16, 3460.

Botha, S., Nass, K., Barends, T. R. M., Kabsch, W., Latz, B., Dworkowski, F., Foucar, L., Panepucci, E., Wang, M., Shoeman, R. L., Schlichting, I. & Doak, R. B. (2015). Acta Cryst. D 71, 387–397.

Chapman, H. N. (2017). Vol. Protein Crystallography: Methods and Protocols, edited by A. Wlodawer, Z. Dauter & M. Jaskolski. pp. 295–324. Humana Press.

Chapman, H. N., Caleman, C. & Timneanu, N. (2014). Philos. Trans. R. Soc. B 369, 20130313.

Crosson, S. & Moffat, K. (2001). Proc. Natl Acad. Sci. USA 98, 2995–3000.

De Zitter, E., Coquelle, N., Oeser, P., Barends, T. R. M. & Colletier, J.-P. (2022). Commun. Biol. 5, 640.

Emsley, P., Lohkamp, B., Scott, W. G. & Cowtan, K. (2010). Acta Cryst. D 66, 486–501.

Fedorov, R., Schlichting, I., Hartmann, E., Domratcheva, T., Fuhrmann, M. & Hegemann, P. (2003). Biophys. J. 84, 2474–2482.

Genick, U. K., Borgstahl, G. E. O., Ng, K., Ren, Z., Pradervand, C., Burke, P. M., Šrajer, V., Teng, T.-Y., Schildkamp, W., McRee, D. E., Moffat, K. & Getzoff, E. D. (1997). Science 275, 1471–1475.

Gevorkov, Y., Yefanov, O., Barty, A., White, T. A., Mariani, V., Brehm, W., Tolstikova, A., Grigat, R.-R. & Chapman, H. N. (2019). Acta Cryst. A 75, 694–704.

Gonzalez, A., Krojer, T., Nan, J., Bjelčić, M., Aggarwal, S., Gorgisyan, I., Milas, M., Eguiraun, M., Casadei, C., Chenchiliyan, M., Jurgilaitis, A., Kroon, D., Ahn, B., Ekström, J. C., Aurelius, O., Lang, D., Ursby, T. & Thunnissen, M. M. G. M. (2025). J. Synchrotron Rad. 32, 779–791.

Gonzalez, A., Milas, M., Bjelčić, M., Chenchiliyan, M., Casadei, C. M., Lang, D., Begum, A., Rollet, K., Panepucci, E., Bento, I., Kapetanaki, S. M., Malmgren, M., Schröder, G. C., Krojer, T., Nan, J., Thunnissen, M. M. G. M. & Ursby, T. (2026). Synchrotron Radiat. News 39, 18–23.

Gotthard, G., Flores-Ibarra, A., Carrillo, M., Kepa, M. W., Mason, T. J., Stegmann, D. P., Olasz, B., Pachota, M., Dworkowski, F., Ozerov, D., Pedrini, B. F., Padeste, C., Beale, J. H. & Nogly, P. (2024). IUCrJ 11, 749–761.

Gotthard, G., Mous, S., Weinert, T., Maia, R. N. A., James, D., Dworkowski, F., Gashi, D., Furrer, A., Ozerov, D., Panepucci, E., Wang, M., Schertler, G. F. X., Heberle, J., Standfuss, J. & Nogly, P. (2024). IUCrJ 11, 792–808.

Grünbein, M. L., Stricker, M., Nass Kovacs, G., Kloos, M., Doak, R. B., Shoeman, R. L., Reinstein, J., Lecler, S., Haacke, S. & Schlichting, I. (2020). Nat. Methods 17, 681–684.

Harper, S. M., Neil, L. C. & Gardner, K. H. (2003). Science 301, 1541–1544.

Henry, L., Berntsson, O., Panman, M. R., Cellini, A., Hughes, A. J., Kosheleva, I., Henning, R. & Westenhoff, S. (2020). Biochemistry 59, 3206–3215.

Huang, K., Merkle, T. & Beck, C. F. (2002). Physiol. Plant. 115, 613–622.

Ihee, H., Rajagopal, S., Šrajer, V., Pahl, R., Anderson, S., Schmidt, M., Schotte, F., Anfinrud, P. A., Wulff, M. & Moffat, K. (2005). Proc. Natl Acad. Sci. USA 102, 7145–7150.

James, D., Weinert, T., Skopintsev, P., Furrer, A., Gashi, D., Tanaka, T., Nango, E., Nogly, P. & Standfuss, J. (2019). J. Vis. Exp. e59087.

Jung, Y. O., Lee, J. H., Kim, J., Schmidt, M., Moffat, K., Šrajer, V. & Ihee, H. (2013). Nat. Chem. 5, 212–220.

Karplus, P. A. & Diederichs, K. (2012). Science 336, 1030–1033.

Kottke, T., Heberle, J., Hehn, D., Dick, B. & Hegemann, P. (2003). Biophys. J. 84, 1192–1201.

Kovalev, K., Tsybrov, F., Alekseev, A., Shevchenko, V., Soloviov, D., Siletsky, S., Bourenkov, G., Agthe, M., Nikolova, M., von Stetten, D., Astashkin, R., Bukhdruker, S., Chizhov, I., Royant, A., Kuzmin, A., Gushchin, I., Rosselli, R., Rodriguez-Valera, F., Ilyinskiy, N., Rogachev, A., Borshchevskiy, V., Schneider, T. R., Bamberg, E. & Gordeliy, V. (2023). Nat. Struct. Mol. Biol. 30, 970–979.

Leonarski, F., Nan, J., Matej, Z., Bertrand, Q., Furrer, A., Gorgisyan, I., Bjelčić, M., Kepa, M., Glover, H., Hinger, V., Eriksson, T., Cehovin, A., Eguiraun, M., Gasparotto, P., Mozzanica, A., Weinert, T., Gonzalez, A., Standfuss, J., Wang, M., Ursby, T. & Dworkowski, F. (2023). IUCrJ 10, 729–737.

Liebschner, D., Afonine, P. V., Baker, M. L., Bunkóczi, G., Chen, V. B., Croll, T. I., Hintze, B., Hung, L.-W., Jain, S., McCoy, A. J., Moriarty, N. W., Oeffner, R. D., Poon, B. K., Prisant, M. G., Read, R. J., Richardson, J. S., Richardson, D. C., Sammito, M. D., Sobolev, O. V., Stockwell, D. H., Terwilliger, T. C., Urzhumtsev, A. G., Videau, L. L., Williams, C. J. & Adams, P. D. (2019). Acta Cryst. D 75, 861–877.

Lomb, L., Barends, T. R. M., Kassemeyer, S., Aquila, A., Epp, S. W., Erk, B., Foucar, L., Hartmann, R., Rudek, B., Rolles, D., Rudenko, A., Shoeman, R. L., Andreasson, J., Bajt, S., Barthelmess, M., Barty, A., Bogan, M. J., Bostedt, C., Bozek, J. D., Caleman, C., Coffee, R., Coppola, N., DePonte, D. P., Doak, R. B., Ekeberg, T., Fleckenstein, H., Fromme, P., Gebhardt, M., Graafsma, H., Gumprecht, L., Hampton, C. Y., Hartmann, A., Hauser, G., Hirsemann, H., Holl, P., Holton, J. M., Hunter, M. S., Kabsch, W., Kimmel, N., Kirian, R. A., Liang, M., Maia, F. R. N. C., Meinhart, A., Marchesini, S., Martin, A. V., Nass, K., Reich, C., Schulz, J., Seibert, M. M., Sierra, R., Soltau, H., Spence, J. C. H., Steinbrener, J., Stellato, F., Stern, S., Timneanu, N., Wang, X., Weidenspointner, G., Weierstall, U., White, T. A., Wunderer, C., Chapman, H. N., Ullrich, J., Strüder, L. & Schlichting, I. (2011). Phys. Rev. B 84, 214111.

Losi, A., Gardner, K. H. & Möglich, A. (2018). Chem. Rev. 118, 10659–10709.

Malla, T. N., Muniyappan, S., Menendez, D., Ogukwe, F., Dale, A. N., Clayton, J. D., Weatherall, D. D., Karki, P., Dangi, S., Mandella, V., Pacheco, A. A., Stojković, E. A., Rose, S. L., Orlans, J., Basu, S., de Sanctis, D. & Schmidt, M. (2025). IUCrJ 12, 36–48.

Martin-Garcia, J. M. (2021). Crystals 11, 521.

McCoy, A. J., Grosse-Kunstleve, R. W., Adams, P. D., Winn, M. D., Storoni, L. C. & Read, R. J. (2007). J. Appl. Cryst. 40, 658–674.

Mehrabi, P., Bücker, R., Bourenkov, G., Ginn, H. M., von Stetten, D., Müller-Werkmeister, H. M., Kuo, A., Morizumi, T., Eger, B. T., Ou, W.-L., Oghbaey, S., Sarracini, A., Besaw, J. E., Paré-Labrosse, O., Meier, S., Schikora, H., Tellkamp, F., Marx, A., Sherrell, D. A., Axford, D., Owen, R. L., Ernst, O. P., Pai, E. F., Schulz, E. C. & Miller, R. J. D. (2021). Sci. Adv. 7, eabf1380.

Mehrabi, P., Müller-Werkmeister, H. M., Leimkohl, J.-P., Schikora, H., Ninkovic, J., Krivokuca, S., Andriček, L., Epp, S. W., Sherrell, D., Owen, R. L., Pearson, A. R., Tellkamp, F., Schulz, E. C. & Miller, R. J. D. (2020). J. Synchrotron Rad. 27, 360–370.

Mehrabi, P., Schulz, E. C., Dsouza, R., Müller-Werkmeister, H. M., Tellkamp, F., Miller, R. J. D. & Pai, E. F. (2019). Science 365, 1167–1170.

Moffat, K. (1989). Annu. Rev. Biophys. Biophys. Chem. 18, 309–332.

Moffat, K. (2001). Chem. Rev. 101, 1569–1582.

Mozzanica, A., Andrä, M., Barten, R., Bergamaschi, A., Chiriotti, S., Brückner, M., Dinapoli, R., Fröjdh, E., Greiffenberg, D., Leonarski, F., Lopez-Cuenca, C., Mezza, D., Redford, S., Ruder, C., Schmitt, B., Shi, X., Thattil, D., Tinti, G., Vetter, S. & Zhang, J. (2018). Synchrotron Rad. News 31, 16–20.

Mueller, U., Thunnissen, M., Nan, J., Eguiraun, M., Bolmsten, F., Milàn-Otero, A., Guijarro, M., Oscarsson, M., de Sanctis, D. & Leonard, G. (2017). Synchrotron Rad. News 30, 22–27.

Nam, K. H. (2024). Curr. Res. Struct. Biol. 7, 100131.

Nass, K., Foucar, L., Barends, T. R. M., Hartmann, E., Botha, S., Shoeman, R. L., Doak, R. B., Alonso-Mori, R., Aquila, A., Bajt, S., Barty, A., Bean, R., Beyerlein, K. R., Bublitz, M., Drachmann, N., Gregersen, J., Jönsson, H. O., Kabsch, W., Kassemeyer, S., Koglin, J. E., Krumrey, M., Mattle, D., Messerschmidt, M., Nissen, P., Reinhard, L., Sitsel, O., Sokaras, D., Williams, G. J., Hau-Riege, S., Timneanu, N., Caleman, C., Chapman, H. N., Boutet, S. & Schlichting, I. (2015). J. Synchrotron Rad. 22, 225–238.

Niziński, S., Marekha, B., Reinstein, J., Shoeman, R. L., Doak, R. B. & Schlichting, I. (2025). J. Appl. Cryst. 58, 2078–2089.

Nogly, P., James, D., Wang, D., White, T. A., Zatsepin, N., Shilova, A., Nelson, G., Liu, H., Johansson, L., Heymann, M., Jaeger, K., Metz, M., Wickstrand, C., Wu, W., Båth, P., Berntsen, P., Oberthuer, D., Panneels, V., Cherezov, V., Chapman, H., Schertler, G., Neutze, R., Spence, J., Moraes, I., Burghammer, M., Standfuss, J. & Weierstall, U. (2015). IUCrJ 2, 168–176.

Okajima, K., Aihara, Y., Takayama, Y., Nakajima, M., Kashojiya, S., Hikima, T., Oroguchi, T., Kobayashi, A., Sekiguchi, Y., Yamamoto, M., Suzuki, T., Nagatani, A., Nakasako, M. & Tokutomi, S. (2014). J. Biol. Chem. 289, 413–422.

Olasz, B., Gotthard, G., Nag, P., González-Viegas, M., Mous, S., Koczurowska, A., Johnson, P. J. M., Sato, T., Shankar, M. K., Wranik, M., Solecka, A., Melo, D., Ke, R., James, D., Nass, K., Langner, P., Valerio, J., Furrer, A., Gashi, D., de Wijn, R., Popelar, T., Pachota, M., Wisniewska, M., Dietze, T., E, J., Asghar, A., Zabelskii, D., Trost, F., Koua, F. H. M., Smyth, P., Sobolev, E., Kim, C., Ozerov, D., Letrun, R., Turkot, O., Doerner, K., Bielecki, J., Han, H., Dworkowski, F., Cirelli, C., Schertler, G. F. X., Schulz, J., Bacellar, C., Standfuss, J., Bean, R., Milne, C., Heberle, J., Schapiro, I. & Nogly, P. (2026). bioRxiv 10.64898/2026.08.02.742302.

Pearson, A. R. & Mehrabi, P. (2020). Curr. Opin. Struct. Biol. 65, 168–174.

Ren, Z., Perman, B., Šrajer, V., Teng, T.-Y., Pradervand, C., Bourgeois, D., Schotte, F., Ursby, T., Kort, R., Wulff, M. & Moffat, K. (2001). Biochemistry 40, 13788–13801.

Salomon, M., Christie, J. M., Knieb, E., Lempert, U. & Briggs, W. R. (2000). Biochemistry 39, 9401–9410.

Schmidt, M. (2008). Vol. Ultrashort Laser Pulses in Biology and Medicine, edited by M. Braun, P. Gilch & W. Zinth. pp. 201–241. Springer.

Schmidt, M. (2013). Adv. Condens. Matter Phys. 2013, 1–10.

Schmidt, M. (2020). Crystals 10, 116.

Schmidt, M. (2023). Vol. Integrated Structural Biology, edited by T. Polenova, C. M. Quinn & A. M. Gronenborn. pp. 203–240. The Royal Society of Chemistry.

Schotte, F., Cho, H. S., Kaila, V. R. I., Kamikubo, H., Dashdorj, N., Henry, E. R., Graber, T. J., Henning, R., Wulff, M., Hummer, G., Kataoka, M. & Anfinrud, P. A. (2012). Proc. Natl Acad. Sci. USA 109, 19256–19261.

Schulz, E. C., Mehrabi, P., Müller-Werkmeister, H. M., Tellkamp, F., Jha, A., Stuart, W., Persch, E., De Gasparo, R., Diederich, F., Pai, E. F. & Miller, R. J. D. (2018). Nat. Methods 15, 901–904.

Shilova, A., Lebrette, H., Aurelius, O., Nan, J., Welin, M., Kovacic, R., Ghosh, S., Safari, C., Friel, R. J., Milas, M., Matej, Z., Högbom, M., Brändén, G., Kloos, M., Shoeman, R. L., Doak, B., Ursby, T., Håkansson, M., Logan, D. T. & Mueller, U. (2020). J. Synchrotron Rad. 27, 1095–1102.

Studier, F. W. (2005). Protein Expr. Purif. 41, 207–234.

Sugahara, M., Nakane, T., Masuda, T., Suzuki, M., Inoue, S., Song, C., Tanaka, R., Nakatsu, T., Mizohata, E., Yumoto, F., Tono, K., Joti, Y., Kameshima, T., Hatsui, T., Yabashi, M., Nureki, O., Numata, K., Nango, E. & Iwata, S. (2017). Sci. Rep. 7, 703.

Swartz, T. E., Corchnoy, S. B., Christie, J. M., Lewis, J. W., Szundi, I., Briggs, W. R. & Bogomolni, R. A. (2001). J. Biol. Chem. 276, 36493–36500.

Weierstall, U., James, D., Wang, C., White, T. A., Wang, D., Liu, W., Spence, J. C. H., Doak, R. B., Nelson, G., Fromme, P., Fromme, R., Grotjohann, I., Kupitz, C., Zatsepin, N. A., Liu, H., Basu, S., Wacker, D., Han, G. W., Katritch, V., Boutet, S., Messerschmidt, M., Williams, G. J., Koglin, J. E., Seibert, M. M., Klinker, M., Gati, C., Shoeman, R. L., Barty, A., Chapman, H. N., Kirian, R. A., Beyerlein, K. R., Stevens, R. C., Li, D., Shah, S. T. A., Howe, N., Caffrey, M. & Cherezov, V. (2014). Nat. Commun. 5, 3309.

Weinert, T., Skopintsev, P., James, D., Dworkowski, F., Panepucci, E., Kekilli, D., Furrer, A., Brünle, S., Mous, S., Ozerov, D., Nogly, P., Wang, M. & Standfuss, J. (2019). Science 365, 61–65.

White, T. A., Kirian, R. A., Martin, A. V., Aquila, A., Nass, K., Barty, A. & Chapman, H. N. (2012). J. Appl. Cryst. 45, 335–341.

White, T. A., Mariani, V., Brehm, W., Yefanov, O., Barty, A., Beyerlein, K. R., Chervinskii, F., Galli, L., Gati, C., Nakane, T., Tolstikova, A., Yamashita, K., Yoon, C. H., Diederichs, K. & Chapman, H. N. (2016). J. Appl. Cryst. 49, 680–689.

Williams, C. J., Headd, J. J., Moriarty, N. W., Prisant, M. G., Videau, L. L., Deis, L. N., Verma, V., Keedy, D. A., Hintze, B. J., Chen, V. B., Jain, S., Lewis, S. M., Arendall, W. B., Snoeyink, J., Adams, P. D., Lovell, S. C., Richardson, J. S. & Richardson, D. C. (2018). Protein Sci. 27, 293–315.

Winn, M. D., Ballard, C. C., Cowtan, K. D., Dodson, E. J., Emsley, P., Evans, P. R., Keegan, R. M., Krissinel, E. B., Leslie, A. G. W., McCoy, A., McNicholas, S. J., Murshudov, G. N., Pannu, N. S., Potterton, E. A., Powell, H. R., Read, R. J., Vagin, A. & Wilson, K. S. (2011). Acta Cryst. D 67, 235–242.

Zeldin, O. B., Gerstel, M. & Garman, E. F. (2013). J. Appl. Cryst. 46, 1225–1230.

Zhang, S., Abiven, Y.-M., Bisou, J., Langlois, F., Renaud, G., Ta, F., Thibaux, G., Abbott, M., Cobb, T., Turner, C., Uzun, I. & Minolli, S. (2018). Proceedings of the 16th Int. Conf. on Accelerator and Large Experimental Control Systems ICALEPCS2017, 143–150.

